# Long-Range Coupling of Posterior Cell Addition and Anterior Vacuolation Provides Robustness in Notochord Elongation

**DOI:** 10.64898/2026.02.17.706348

**Authors:** Carlos Camacho-Macorra, Alberto Ceccarelli, Dillan Saunders, Guillermo Serrano Nájera, Osvaldo Chara, Benjamin Steventon

**Affiliations:** Department of Genetics, University of Cambridge, Cambridge, UK, CB2 3EH; Selwyn College, University of Cambridge, UK, CB3 9DQ; School of Biosciences, University of Nottingham, UK, LE12 5RD; Instituto de Tecnología, Universidad Argentina de la Empresa, Buenos Aires, Argentina

## Abstract

Robust tissue growth control requires long-range communication between the rate of progenitor addition and tissue expansion. However, the regulatory mechanisms that couple these two processes are unknown. In zebrafish, notochord morphogenesis is a principal driver of axis extension through the combined actions of posterior progenitor addition and anterior vacuolation. To elucidate how progenitor dynamics and vacuole-driven cell expansion interact to shape notochord development, we first generated a mathematical model that links progenitor addition rate to the progressive expansion of cells from anterior-to-posterior to simulate vacuolation rate. Comparing this with empirical measurements, we find that progenitor incorporation together with vacuolation, produces a linear gradient in nearest neighbour distance. We next explored the role of YAP/TAZ in regulating the rate of progenitor addition in mutants for YAP/TAZ inhibitor *vgll4b*. We find that *vgll4b* expression and YAP activity are enriched in posterior midline progenitors. Loss of *vgll4b* elevates YAP signaling, enhances progenitor addition, restricts vacuole expansion, and—after a transient buffering phase—compromises A-P axis elongation. These results support a long-range feedback mechanism linking progenitor recruitment to vacuolation, enabling the notochord to balance cellular input with volumetric expansion, thereby maintaining tissue proportions.

**Graphical abstract. Proposed working model:** 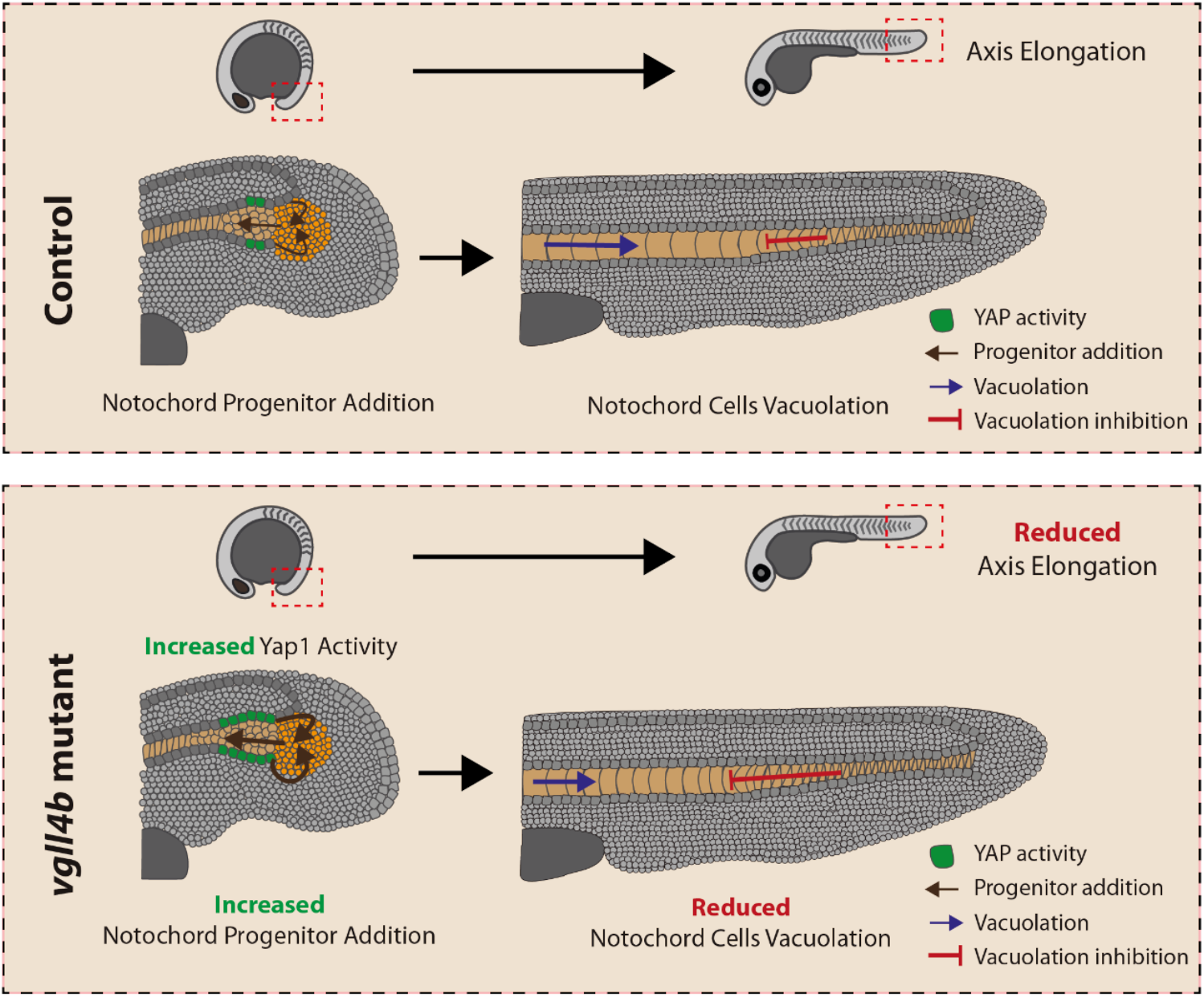

**A**. Schematic representation of the authors’ proposed model illustrating how increased YAP activation in notochord progenitors of *vgll4b* mutants leads to enhanced progenitor addition to the notochord. This increased incorporation subsequently compromises the ability of notochord cells to undergo proper vacuolation, resulting in reduced axial elongation compared with control embryos.

## Introduction

A defining feature of vertebrate development is the elongation of the body along the anterior–posterior (A–P) axis, a process in which the generation of new tissue at the posterior end is coordinated with differentiation and growth in more anterior regions (Bénazéraf et al., 2017; Mongera et al., 2019; Steventon et al., 2016). This process requires the integrated action of cellular, molecular, and mechanical mechanisms across spatially distinct regions of the embryo to maintain proportionate axis elongation. Multiple, partially overlapping, mechanisms contribute to vertebrate axis extension. During gastrulation and early somitogenesis, convergent extension movements narrow and lengthen the embryonic body plan (Keller, 2002; Steventon et al., 2016). Subsequently, distinct populations of progenitors in the tailbud supply cells to the forming neural tube, notochord, and somites (Wymeersch et al., 2021). In parallel, mechanical forces arising from tissue tension, extracellular matrix remodeling, and hydrostatic pressure shape and elongate axial structures (Adams et al., 1990; Irvine and Shraiman, 2017; Mongera et al., 2018). Furthermore, mechanical and spatial interactions between adjacent tissues regulate tissue proportions through a phenomenon referred to as multi-tissue tectonics (Blanchard et al., 2009; Busby and Steventon, 2021; Saunders et al., 2025). Because these processes occur concurrently and mutually influence one another, embryos must coordinate them to ensure robust patterning and preserve axial proportions (Saiz and Hadjantonakis, 2020). Despite extensive work on the cellular sources and molecular signals that govern posterior axis formation, the mechanisms by which embryos balance posterior tissue generation with anterior tissue expansion remain incompletely understood.

This mechanism is particularly relevant for structures such as the zebrafish notochord, a defining feature of the chordate body plan. In addition to providing physical support and patterning cues to surrounding tissues (Halpern, 1997; Stemple, 2005), the notochord plays a central role in mechanically elongating the A–P axis (Adams et al., 1990; McLaren and Steventon, 2021). In zebrafish, midline progenitor cells (MPCs) located in the basal region of the tailbud surrounding the chordoneural hinge (CNH) give rise to the mesodermal notochord, neural floor plate, and endodermal hypochord lineages (Row et al., 2016). The hypochord is a transient endodermal structure present in teleosts and some other vertebrates, positioned ventral to the notochord and contributing to the patterning of surrounding tissues (Hogan and Bautch, 2004).

Zebrafish notochord morphogenesis can be thought of in terms of three distinct phases (Figure 1A). In the first phase, posterior progenitor addition drives elongation by incorporating new cells into the posterior notochord. In the second phase, progenitor addition occurs together with the anterior vacuolation causes differentiated notochord cells to increase in volume, creating a “vacuolation front” that progresses from anterior to posterior along the axis. The resulting pressure generates mechanical forces that stretch and elongate the notochord (Ellis et al., 2013; McLaren and Steventon, 2021; Stemple, 2005). Upon progenitor depletion, a final phase is characterized by continued notochord vacuolation at post-somitogenesis stages of development. Achieving the correct balance between these opposing dynamics is essential for producing a properly proportioned notochord and, by extension, a correctly scaled vertebrate body axis (McLaren and Steventon, 2021). Disruption of either progenitor addition (Talbot et al., 1995) or notochord cell vacuolation (Bagwell et al., 2020; Ellis et al., 2013) results in a shortened A–P axis. Progenitor addition is a key determinant of notochord elongation and cell number, yet insights into the rate of incorporation and the molecular pathways regulating the process remain limited. Understanding how progenitor recruitment is coordinated with tissue-scale processes, such as anterior vacuolation, is essential to elucidating the mechanisms that ensure robust notochord elongation and proper A–P scaling in the zebrafish embryo.

**Figure 1.**
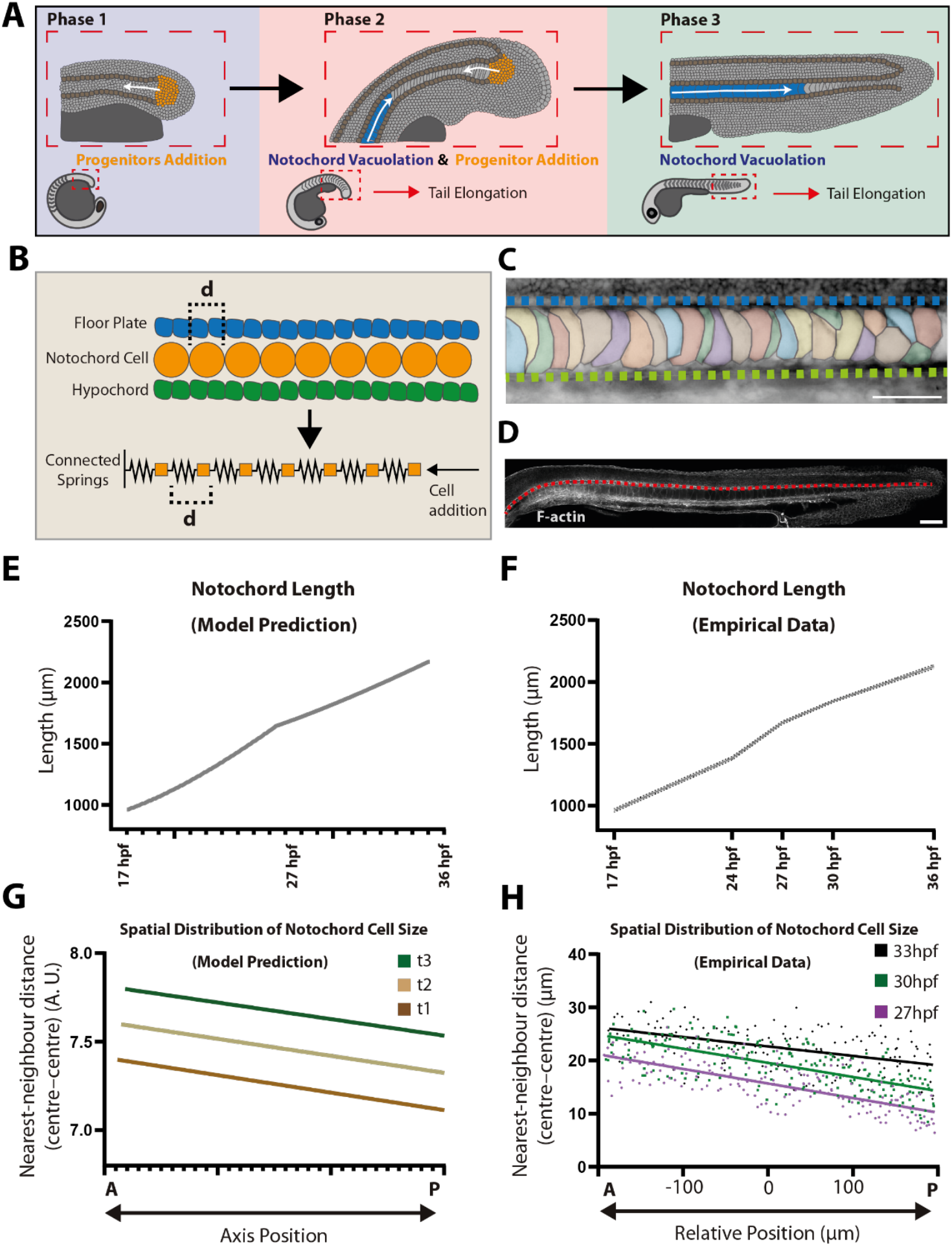
Mathematical model predicting notochord elongation dynamics and scaling properties in zebrafish embryos. **A**. Scheme representing the three phases of late notochord morphogenesis, where the last two are described in the mathematical model. **B**. Scheme illustrating key concepts used to construct the mathematical model. **C**. Examples of notochord vacuole segmentation visualized with Bodipy staining; the blue line marks the notochord, and the green line marks the hypochord (scale bar: 50 μm). **D**. Example of the methodology used to measure notochord length by tracing the floor plate (red line) in confocal images of phalloidin-stained embryos (scale bar: 100 μm). **E**. Mathematical model prediction of notochord length at different time points. **F**. Quantification of A–P axis length at 17 hpf (n = 12), 24 hpf (n = 10), 27 hpf (n = 20), 30 hpf (n = 16), and 36 hpf (n = 7). **G**. Mathematical model prediction of AP profiles of centre-to-centre nearest-neighbour cell distances within the notochord at different time points. **H**. Quantification of centre-to-centre nearest-neighbour cell distances within the notochord at 27 hpf, 30 hpf, and 33 hpf. The A–P axis in the graph is centred at the yolk-level extension. The region of interest (ROI) used for measurements spanned 200 µm on each side of this reference point, for a total length of 400 µm.

The Hippo pathway effector YAP, through its interaction with TEAD transcription factors, has emerged as a central regulator of mechanotransduction (Dupont et al., 2011), progenitor maintenance (Han et al., 2015; Wang et al., 2016), and axial development (Porazinski et al., 2015) across vertebrate species. YAP expression has been documented in the notochord across multiple vertebrate models, including mouse (Sawada et al., 2008), frog (Nejigane et al., 2010), and zebrafish (Jiang et al., 2009), indicating an evolutionarily conserved presence in this axial structure. Recent cross-species studies further suggest that YAP plays a functional role in notochord development. In human trunk organoids, YAP contributes to notochord specification (Rito et al., 2025), while in mouse embryos, YAP responds to mechanical inputs to regulate FoxA2 and Shh expression, thereby coordinating notochord and floor plate formation (Cheng et al., 2023). In medaka, YAP drives a mechanosensitive program that sustains collective cell migration required for axis assembly (Sousa-Ortega et al., 2023). In zebrafish, *yap1* and its paralog *wwtr1* are expressed and active in the notochord (Astone et al., 2024; Kimelman et al., 2017) and contribute to posterior body elongation and overall morphogenesis (Kimelman et al., 2017). Despite these insights, the specific roles of YAP in the morphogenesis of the zebrafish notochord and the mechanisms regulating its activity in this context remain incompletely defined.

VGLL4 is a well-established inhibitor of YAP activity (Tang et al., 2024; Zhu et al., 2025), acting through competitive binding to TEAD transcription factors (Deng and Fang, 2018; Shi et al., 2017; Zhang et al., 2021). This antagonism has led to the view that a major function of YAP in development is to counteract a default VGLL4-mediated repression (Cai et al., 2022). During embryogenesis, the VGLL4–TEAD axis regulates diverse processes, including cardiac development (Lin et al., 2016; Lin et al., 2016; Sheldon et al., 2022; Yu et al., 2019), skeletal muscle formation (Feng et al., 2019), left–right asymmetry (Fillatre et al., 2019), osteoblast differentiation (Suo et al., 2020), and posterior lateral line primordium migration (Lardennois et al., 2025). However, a potential role for *VGLL4* in notochord morphogenesis has not yet been investigated.

To explore this possibility, we first developed a mathematical model integrating posterior progenitor addition and anterior vacuolation to investigate their combined effects on notochord elongation and cell size distribution. This computational framework allows us to explore how variations in progenitor incorporation rates or vacuolation dynamics influence tissue growth and morphogenesis. Importantly, the model provides a mechanistic basis to test our central hypothesis: that the balance between progenitor addition and vacuolation is actively regulated. By combining *in vivo* measurements, functional manipulations of YAP activity, and computational predictions, we aim to uncover the regulatory principles linking YAP signaling, progenitor behavior, and tissue-scale dynamics during vertebrate axis elongation.

## Results

### Mathematical modelling reveals how posterior progenitor addition and anterior vacuolation drive notochord elongation and scaling of cell sizes

To investigate how posterior progenitor addition and cell vacuolation are coordinated during notochord elongation, we developed a minimal mathematical model representing the notochord as a linear sequence of mass–spring cells along the A-P axis, *x*. Similar biophysical and computational approaches have been used previously to study notochord elongation, cell packing, and morphogenetic mechanics in vertebrate embryos (Adams et al., 1990; Curcio and Lubkin, 2023; Yasuoka, 2020). In our model, progenitor cells are added with a rate *r*_*p*_ at the posterior end at defined intervals by attaching to the terminal cell, while a vacuolation front advances posteriorly at constant speed *v*_front_, triggering linear growth (at a rate *J*) of target cell size *S*_*i*_ once cells lie anterior to the front (Figure 1B-D). To better understand the coordination of these processes, we focus on the last two phases of notochord elongation: when progenitor addition coincides with vacuolation, and upon progenitor depletion when only vacuolation proceeds (Figure 1A). A complete description of the model is provided in the Methods section.

As expected, the model predicts that both cell addition and vacuolation drive notochord elongation. Notably, the modelled notochord elongates approximately linearly in time during both phases, with comparable growth rates (Figure 1E). To test this prediction, we measured notochord length in embryos at multiple time points between 17 and 36 hours post fertilization (hpf) and observed a similar linear growth pattern (Figure 1F). As vacuolation begins anteriorly, cell sizes decrease linearly from anterior to posterior. Over time, this linear profile maintains its negative slope constant while its positive intercept increases (Figure 1G). We refer to this phenotype as scaling, as the overall shape of the distribution is preserved during tissue growth. To test this prediction, we imaged and segmented notochords spanning a 200 μm domain anterior and posterior to the yolk extension from 27 to 33 hpf and quantified the centre-to-centre nearest-neighbour cell distances of notochord cells as a proxy for their cell sizes. Strikingly, the spatial profile of cell sizes along the AP axis is also linear, with a positive intercept that increases over time and a negative slope that remains nearly constant (Figure 1H). Given that, in the zebrafish embryo, a single large fluid-filled vacuole occupies approximately 70–90% of the volume of notochord cells (Bagwell et al., 2020; Ellis et al., 2013), these findings raise the question of how the rate of progenitor addition is coordinated with vacuolation during notochord elongation.

### Active YAP and its inhibitor *vgll4b* co-localize with midline progenitors in the zebrafish tailbud

The zebrafish tailbud contains distinct midline progenitor populations that give rise to mesodermal notochord, neural floor plate, and endodermal hypochord lineages (Latimer and Appel, 2006; Row et al., 2016). Recent work has identified two previously uncharacterized midline progenitor pools located at the most posterior ends of the axial midline (Morabito et al., 2025; Row et al., 2016): a dorsal population at the posterior floor plate that generates both floor plate and notochord, and a ventral population at the posterior hypochord that contributes to hypochord and notochord (Figure 2A). During notochord morphogenesis, these progenitors migrate posteriorly, and cells in the posterior-most region converge into a defined axial progenitor niche to form the extending notochord. In contrast, more anterior trailing progenitors gradually decelerate and differentiate into floor plate or hypochord, respectively.

**Figure 2.**
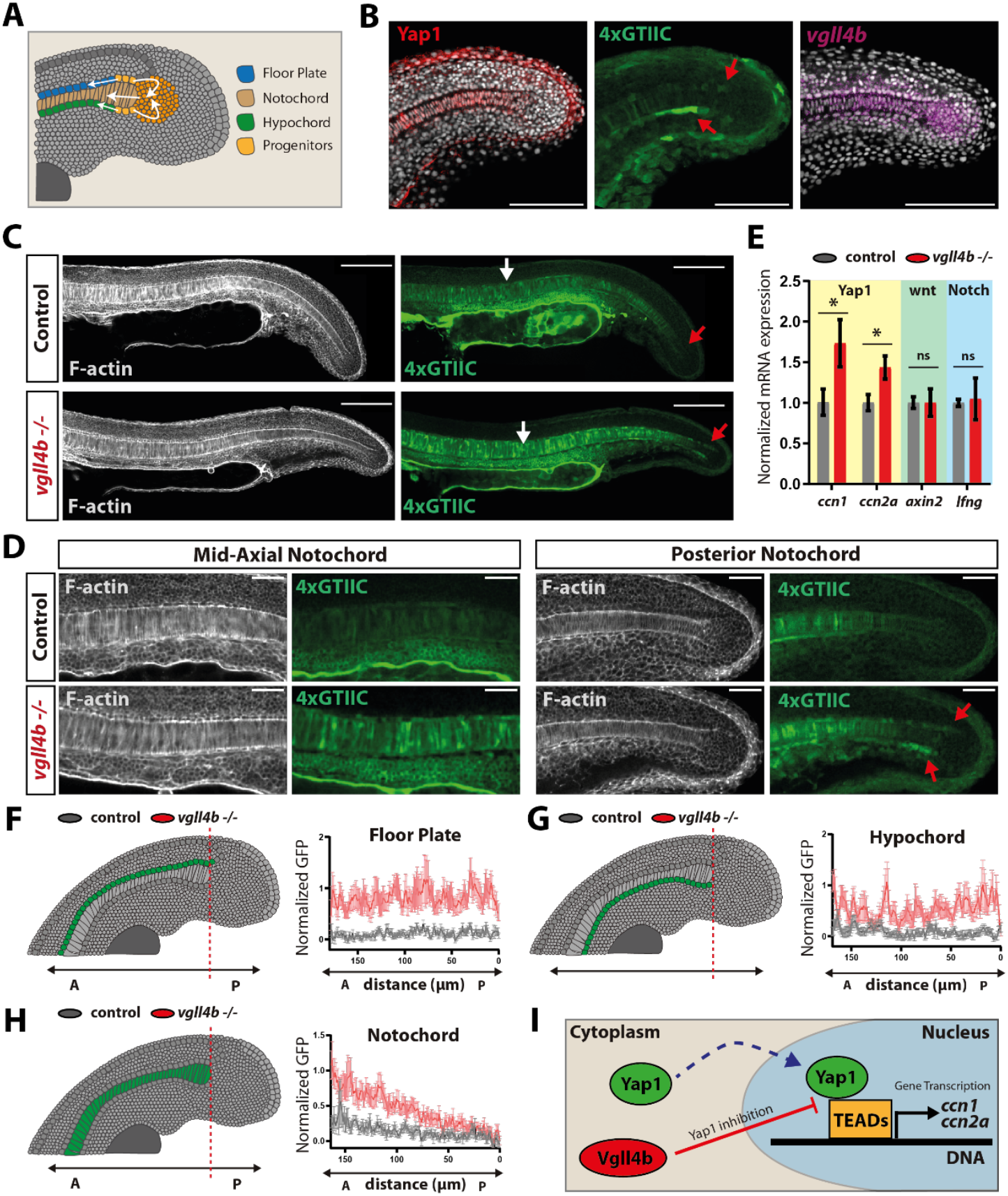
YAP activity is observed in midline progenitors during axis elongation, and it is upregulated in *vgll4b* mutants. **A**. Scheme representing different midline progenitor populations in the zebrafish tailbud. **B**. Confocal images of 24 hpf *Tg(4xGTIIC:GFP)* embryos with immunostaining for YAP and GFP, and HCR for *vgll4b* mRNA (scale bar: 100 μm). Red arrows point GFP expression in the posterior floor plate and hypochord. **C**. Confocal images of 24 hpf *Tg(4xGTIIC:GFP)* control and *Tg(4xGTIIC:GFP)*; *vgll4b*−/− embryos. Red arrows point GFP expression in the progenitor’s region and white arrows point GFP expression in the anterior notochord (scale bar: 100 μm). **D**. Confocal images of the mid-axial and posterior notochord in control and mutant embryos at 24 hpf (scale bar: 50 μm), red arrows point increased GFP expression in the mutant. **E**. qPCR analysis of 24 hpf tailbuds showing upregulation of YAP targets *ccn1* (t-test, p = 0.0305) and *ccn2a* (t-test, p = 0.0157), while *axin2* (t-test, p = 0.9985) and *lfng* (t-test, p = 0.7785) remain unchanged. **F, G, H**. Quantification of normalized GFP intensity across the floor plate, notochord, and hypochord at 24 hpf, showing increased reporter activity in mutants (control n = 13; mutant n = 15). **I**. Scheme illustrating the Vgll4b–YAP inhibition mechanism.

To investigate factors that may regulate the rate at which notochord progenitors are incorporated into the posterior notochord, we focused on YAP, a mechanotransduction effector increasingly recognized as a key regulator of axial development (Sousa-Ortega et al., 2023). Using the YAP activity reporter line *4xGTIIC:GFP*, which marks cells with active YAP signalling (Miesfeld and Link, 2014), together with Yap1 immunostaining, we assessed Yap1 protein localization and activity in the tail during notochord morphogenesis. Consistent with previous findings (Kimelman et al., 2017), Yap1 protein was detected in the epidermis and in midline structures, including the notochord, floor plate, and hypochord (Figure 2B). Reporter activity highlighted strong YAP activation at the posterior ends of the floor plate and hypochord, whereas more anterior regions showed clear activity also in the notochord (Figure 2C, 2D). We next examined the gene expression of zebrafish VGLL4 paralogues during posterior axis elongation and found that *vgll4b* is the only paralogue expressed in the posterior tailbud (Supplementary Figure 1). Its expression is localized to midline tissues, including the notochord, floor plate, and hypochord, as well as to the region containing the earliest described notochord progenitors (Figure 2B).

In summary, these observations demonstrate that YAP is actively engaged within the midline progenitor domain during zebrafish axis elongation, and that its inhibitor *vgll4b* is specifically expressed in the same axial tissues. This spatial overlap indicates that *vgll4b* could modulate YAP activity *in situ*, providing a mechanistic framework for how YAP signalling may regulate progenitor allocation and behaviour during notochord formation.

### *vgll4b* is required to limit YAP activity in the midline progenitors

To determine whether *vgll4b* regulates YAP activity during A-P axis elongation, we used a *vgll4b* loss-of-function line generated into the *4xGTIIC:GFP* reporter background (Camacho-Macorra et al., 2024). Comparing GFP reporter expression in control and mutant embryos revealed a general increase in midline YAP activity in *vgll4b* mutants (Figure 2C). Higher-resolution analysis showed elevated GFP signal in the mid-axial notochord (quantified in Supplementary Figure 2) and in the floor plate. Strikingly, at the posterior end of the axis, YAP overactivation was most prominent in the posterior floor plate and posterior hypochord, corresponding to the region where notochord progenitors reside (Figure 2D).

qPCR on isolated tailbuds from control and mutant embryos showed that YAP transcriptional targets *ccn1* and *ccn2a* were upregulated in *vgll4b* mutants, whereas transcriptional readouts of other signalling pathways known to act in tailbud progenitors were unchanged (Figure 2E). We quantified GFP intensity at 24 hpf across the posterior ends of the three midline tissues—the floor plate (Figure 2F), hypochord (Figure 2G), and notochord (Figure 2H). In all three, YAP reporter activity was consistently higher in *vgll4b* mutants. Similar results were observed at later developmental stages (Supplementary Figure 2).

Together, these findings demonstrate that *vgll4b* mutants exhibit YAP overactivation throughout the posterior midline, including the notochord progenitor niche. This raises the question of whether elevated YAP activity in the mutant perturbs the rate or pattern of progenitor recruitment during notochord formation and posterior axis elongation.

### Loss of *vgll4b* expression enhances notochord progenitor addition

To investigate the requirement of *vgll4b* in notochord progenitor addition, we performed a lineage tracing experiment using the photoconvertible protein Kikume Green-Red (KikGR). Embryos were injected at the one-cell stage with *KikGR* mRNA, and at the onset of tail elongation, a vertical band of cells was photoconverted, encompassing cells destined to form part of the spinal cord, midline structures, and somitic mesoderm (Figure 3A).

**Figure 3.**
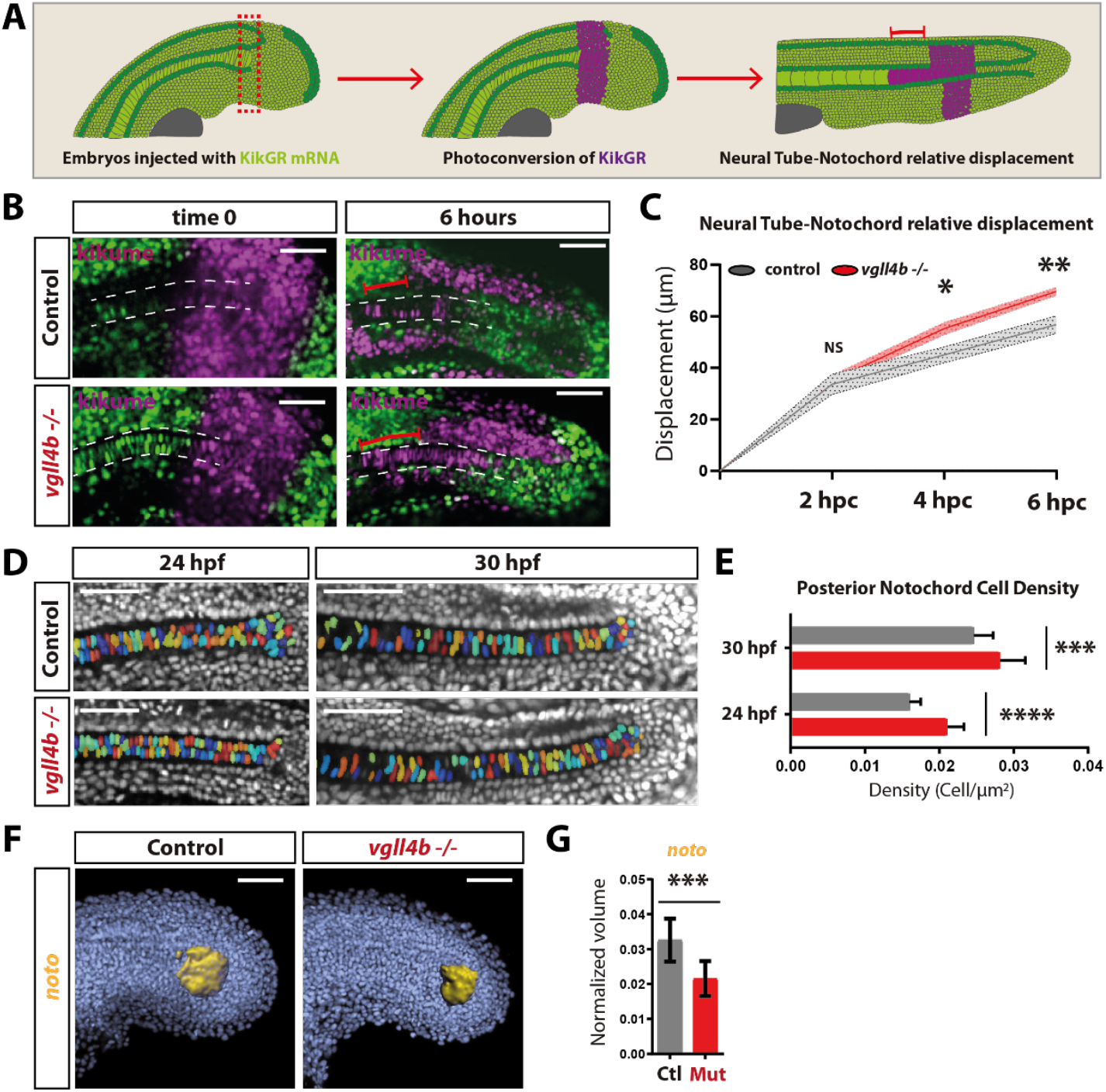
Enhanced YAP activity in *vgll4b* mutants increases the contribution of midline progenitors to the notochord. **A**. Scheme showing the KikGR photoconversion strategy. **B**. Confocal images of photoconverted control and mutant embryos at different time points (scale bar: 100 μm). **C**. Quantification of relative notochord displacement with respect to the neural tube at 2 hpc (t-test; control n = 4, mutant n = 5; p = 0.7288), 4 hpc (t-test; control n = 5, mutant n = 5; p = 0.0306), and 6 hpc (t-test; control n = 4, mutant n = 5; p = 0.0079). **D**. Confocal images of the posterior notochord at 24 hpf and 30 hpf in *Tg(4xGTIIC:GFP)* control and *Tg(4xGTIIC:GFP)*; *vgll4b*−/− embryos, showing segmented nuclei (scale bar: 50 μm). **E**. Quantification of cell density at the posterior end of the notochord at 24 hpf (t-test; control n = 12, mutant n = 18; p < 0.0001) and 30 hpf (t-test; control n = 19, mutant n = 23; p = 0.0008). **F**. Volumes of *noto* expression from HCR staining in control and mutant embryos at 24 hpf (scale bar: 100 μm). **G**. Quantification of normalized *noto* expression volume (t-test; control n = 12, mutant n = 10; p = 0.0004).

If YAP overactivation in the *vgll4b* mutant specifically affects midline progenitor behavior rather than the general addition of cells to the axis, the relative displacement over time between the most anterior photoconverted cells in the notochord and the most anterior photoconverted cells in the spinal cord can serve as a proxy for progenitor addition to the notochord. Consistent with this hypothesis, we observed that in *vgll4b* mutants, the relative displacement between the anterior fronts of photoconverted notochord and spinal cord cells increased over time (Figure 3 B-C), indicating enhanced incorporation of cells into the notochord. To corroborate this finding, we quantified cell density at the posterior end of the notochord at 24 and 30 hpf in control and *vgll4b* mutant embryos (Figure 3D). At both time points, the posterior notochord of *vgll4b* mutants contained more cells than that of controls (Figure 3E), further supporting the hypothesis of increased progenitor addition in the mutants.

To characterise the classical notochord progenitor niche, which is identified by *noto* expression, we measured the volume of the normalized *noto*-expressing domain at 24 hpf (Figure 3F). The *vgll4b* mutants exhibited a reduction in the *noto*-expressing volume at this stage (Figure 3G), suggesting faster depletion of this progenitor pool, consistent with the observed enhanced addition of progenitors into the notochord. To directly assess the increased incorporation of cells into the notochord, we quantified nuclear number and distribution in the mid-axial notochord at the level of the yolk extension (Supplementary Figure 3 A-B). *vgll4b* mutants exhibited a higher number of notochord cells (Supplementary Figure 3C) as well as altered nuclear positioning (Supplementary Figure 3D). In controls, nuclei were primarily localized near the periphery of the notochord, whereas in mutants, the increased cell number resulted in nuclei also occupying more central positions within the structure. To exclude the possibility that mutant embryos contain a higher number of notochord cells prior to tail formation, we compared notochord cell density and total notochord cell number at 17 hpf and found no differences between control and vgll4b mutant embryos in either measure (Supplementary Figure 3 E-G).

Taken together, these observations demonstrate that *vgll4b* mutants, in which YAP activity is elevated in midline structures, exhibit a specific increase in the addition of progenitor cells to the notochord compared with control embryos.

### Feedback between progenitor addition and vacuolation controls notochord morphogenesis

To determine the effects of enhanced posterior progenitor addition on notochord elongation rate, we quantified notochord length at multiple stages from the onset of tail elongation to 36 hpf in control and *vgll4b* mutant embryos (Figure 4A). These measurements reveal a robust size-buffering property during the early phase of notochord extension, when posterior progenitor addition is the predominant driver of elongation. During this period, notochord length remains comparable between genotypes despite the elevated progenitor incorporation observed in *vgll4b* mutants. This buffering later fails in the subsequent phase, when vacuolation becomes the principal driver of elongation (Figure 4A). Here, *vgll4b* mutants display a shorter notochord even though they contribute more progenitor cells to the tissue. This phenotype suggests the presence of a negative feedback mechanism linking posterior progenitor addition to anterior vacuolation.

**Figure 4.**
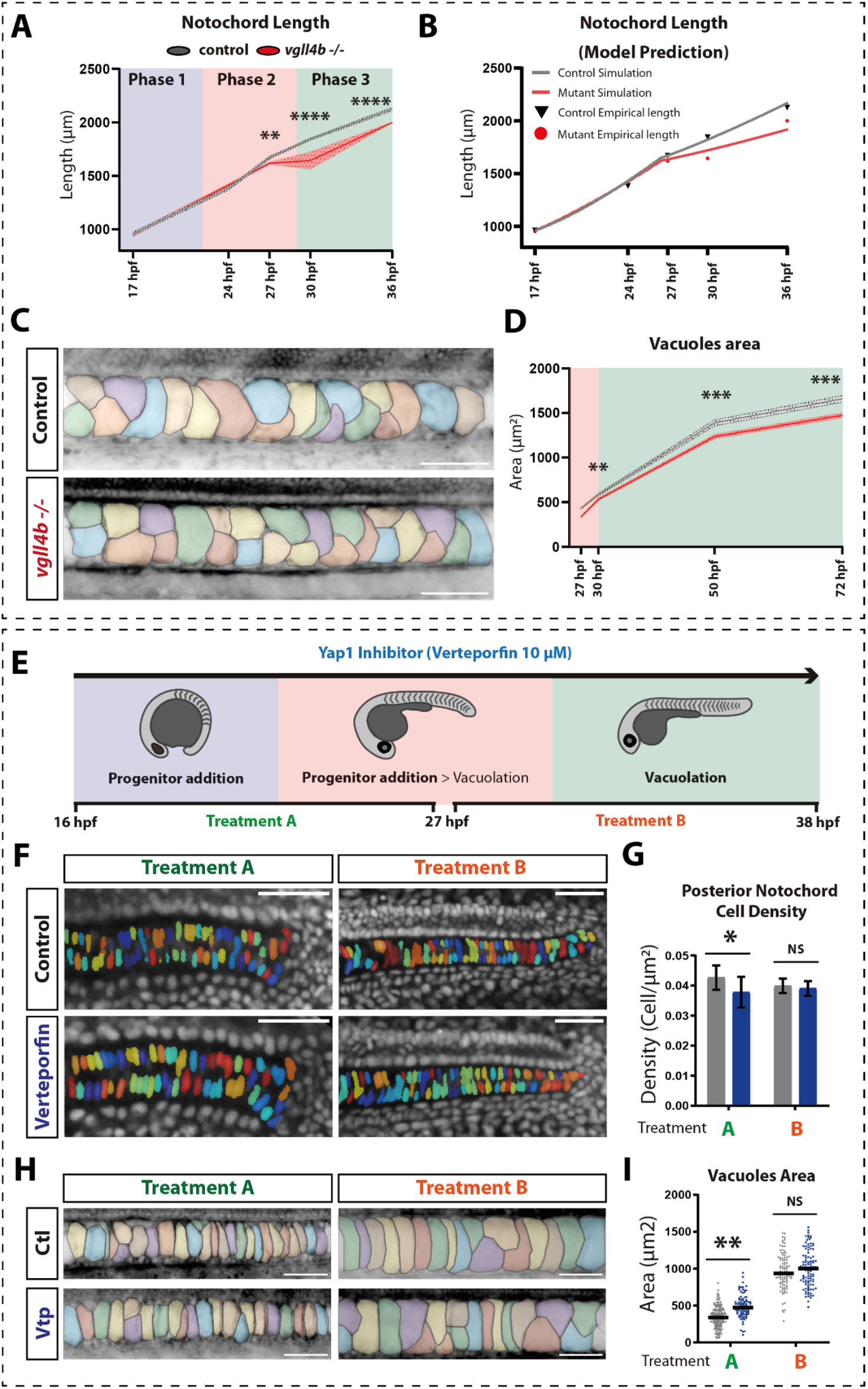
*vgll4b* mutants exhibit notochord vacuolation defects and reduced notochord length, while YAP inhibition decreases notochord cell density and disrupts vacuolation. **A**. Quantification of A–P axis length at 17 hpf (t-test; control n = 12, mutant n = 12; p = 0.7367), 24 hpf (t-test; control n = 10, mutant n = 11; p = 0.1232), 27 hpf (t-test; control n = 20, mutant n = 20; p = 0.0039), 30 hpf (Mann–Whitney test; control n = 16, mutant n = 19; p < 0.0001), and 36 hpf (t-test; control n = 7, mutant n = 7; p < 0.0001). The same control measurements were used in Figure 1F. **B**. Mathematical model predictions of notochord length for control and *vgll4b* mutant embryos at different time points. The model assumes that YAP increases progenitor addition *r*_*p*_ and simultaneously regulates the vacuolation front speed *v*_front_, slowing the wave of vacuolation, without affecting the vacuolation rate of each notochord cell (see Eqs. 3 and 4 in the main text). The empirical measurements shown in the graph are the same as those presented in Figure 4A. **C**. Segmentation of notochord vacuoles in control and mutant embryos at 30 hpf visualized with Bodipy staining (scale bar: 50 μm). **D**. Quantification of notochord vacuole area at 27 hpf (Mann–Whitney test; control n = 8, mutant n = 6; p < 0.0001), 30 hpf (t-test; control n = 8, mutant n = 7; p = 0.0083), 2 dpf (t-test; control n = 9, mutant n = 12; p = 0.0003), and 3 dpf (t-test; control n = 11, mutant n = 12; p = 0.0009). **E**. Schematic of verteporfin (YAP inhibitor) treatment strategy. **F**. Confocal images of the posterior notochord at 24 hpf and 30 hpf in control and verteporfin-treated embryos, showing segmented nuclei (scale bar: 50 μm). **G**. Quantification of posterior notochord cell density: Treatment A (t-test; control n = 10, treated n = 14; p = 0.0219); Treatment B (t-test; control n = 8, treated n = 9; p = 0.4696). H. Segmentation of notochord vacuoles in control and verteporfin-treated embryos visualized with Bodipy staining (scale bar: 50 μm). **I**. Quantification of vacuole area in the central notochord: Treatment A (t-test; control n = 7, treated n = 4; p = 0.00078); Treatment B (t-test; control n = 7, treated n = 7; p = 0.2406).

To investigate this hypothesis, we incorporated it into the mathematical model. We posited that YAP activity negatively regulates vacuolation-induced volume expansion while positively regulating progenitor addition. These effects were incorporated by expressing the vacuolation rate *J* and progenitor addition rate *r*_*p*_ as functions of normalized YAP levels *Y*:

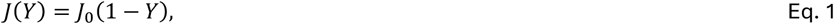

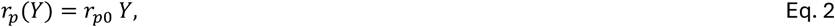

where *J*_0_ and *r*_*p*0_ denote the maximal vacuolation and progenitor addition rates, respectively. Further justification for these functional forms is provided in the Methods.

To investigate the plausibility of this hypothetical feedback between YAP and vacuolation, we analysed experimental AP profiles of nearest-neighbour cell distances for both wild-type and mutant embryos (Figure 1H; Supplementary Figure 4). Assuming scaling (see Methods), we estimated the parameters *J*and *v*_front_ by fitting a linear function to these profiles (see Methods). Under this hypothesis, we expected the vacuolation rate *J* to be reduced in the mutant, while the vacuolation front speed *v*_front_ would remain unchanged. Such a model would be consistent with a mechanism by which YAP activity in the middle region of the notochord (Figure 2C; white arrow) inhibits vacuolation rate, in addition to its role in promoting progenitor addition. However, our estimates yielded *J* = 0.66 ± 0.07 and *v*_front_ = 70 ± 10 for the wild type, and *J* = 0.9 ± 0.2 and *v*_front_ = 110 ± 70 for the mutant. Furthermore, within the experimentally constrained ranges of *J* and *v*_front_, the model failed to reproduce the notochord elongation dynamics, in particular the phase 3 behaviour in which YAP-dependent effects on vacuolation are observed following an earlier phase where these effects are buffered by progenitor addition (Supplementary Figure 5). We therefore rejected the hypothesis that YAP modulates both the progenitor addition rate and the vacuolation rate.

We next proposed an alternative model variant in which YAP regulates the vacuolation front speed *v*_front_. This model would be consistent with an alternate mechanism by which the timing of vacuolation onset is slowed in *vgll4b* mutants, slowing the wave of vacuolation through the tissue. In this formulation, the vacuolation rate *J* is independent of YAP, and thus identical in wild-type and mutant embryos, while, as in the previous model variant, the progenitor addition rate remains higher in the mutant than in the wild type. This revised model is described by the following Eqs. 3 and 4.

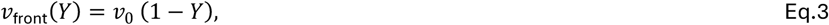

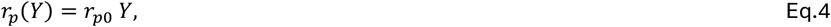

Where *v*_0_ is a constant.

By adopting values of *J* and *v*_front_ within the experimentally derived constraints, we identified a combination that reproduced the notochord elongation dynamics in agreement with that observed experimentally (Figure 4B). This model assumes that in vgll4b mutants, YAP increases the rate of progenitor addition with an additional decrease in the vacuolation front speed, without affecting the vacuolation rate. To test this hypothetic alternative experimentally, we next examined vacuolation dynamics directly in *vgll4b* mutants. Notochord vacuoles were segmented and quantified in the mid-axial notochord from 27 to 72 hpf (Figure 4C). Across all time points, *vgll4b* mutants consistently displayed reduced vacuole areas relative to controls (Figure 4D). This reduction aligns with our earlier observation that mutants contain a greater number of notochord cells, which likely inhibits the ability of a cell to vacuolate. Impaired vacuolation results in a shorter notochord (Figure 4A) and, consequently, a reduced body axis length at 2–3 dpf (Supplementary Figure 6). Shortened axial length is a well-documented outcome in zebrafish embryos with defective notochord vacuolation (Parsons et al., 2002) or fragmented vacuoles (Bagwell et al., 2020; Garcia et al., 2017; Lim et al., 2017). Notably, *vgll4b* mutants also lose the characteristic anterior–posterior scaling of notochord cell A-P length observed in controls (Figure 1H; Supplementary Figure 4), suggesting that altered progenitor addition dynamics disrupt the intrinsic scaling properties of the system.

Taken together, these results demonstrate that the notochord integrates a feedback mechanism linking progenitor addition and anterior vacuolation in zebrafish. We propose that elevated YAP activity, by increasing progenitor density in the pre-vacuolated notochord, reduces notochord vacuolation capacity. At early stages, this explains the buffering capacity of the notochord to maintain the same rate of elongation in *vgll4b* mutants, even though progenitor addition is increased. However, upon the cessation of progenitor addition, the inability to propagate vacuolation across the full extent of the notochord results in a decrease in elongation rate.

### YAP inhibition is required for progenitor addition but not for directly inhibiting anterior vacuolation

To test our prosed model, we next aimed to inhibit YAP signalling during a later phase when notochord elongation is driven only by vacuolation. If YAP plays a direct role in inhibiting vacuolation rate, this should still block anterior expansion as we still observe YAP activity in the notochord at post-somitogenesis stages (Supplementary Figure 2A-B). Conversely, if our model is correct, and the observed inhibition in *vgll4b* mutant embryos is an indirect consequence of increased progenitor addition, no adverse effects of vacuolation would be expected at these stages. To achieve temporal control over YAP inhibition, we used the well-established YAP inhibitor Verteporfin to enable temporal control over YAP inhibition during distinct phases of its elongation (Fillatre et al., 2019; Ren et al., 2021; Ye et al., 2020). We designed two treatment windows (Figure 4E). Treatment A consisted of Verteporfin exposure from 16 to 27 hpf—corresponding to the phase in which posterior progenitor addition is ongoing and constitutes the primary driver of notochord and axis elongation. Treatment B consisted of drug exposure from 27 to 38 hpf, a period during which progenitor addition has ceased and vacuolation becomes the main driver of notochord extension.

We first quantified cell density at the posterior end of the notochord using nuclear labelling (Figure 4F). Embryos subjected to Treatment A showed a marked reduction in posterior cell density, whereas Treatment B embryos were indistinguishable from controls (Figure 4G). This result indicates that the number of cells incorporated into posterior notochord is reduced specifically when YAP is inhibited during the progenitor-addition window.

We next quantified vacuolation under both treatment regimes. Notochord vacuoles were segmented and their area measured in the mid-axial notochord (Figure 4H). Treatment A embryos exhibited a significant increase in vacuole area compared with controls, whereas Treatment B embryos did not show detectable differences (Figure 4I). These findings are consistent with a model in which reduced progenitor addition in Treatment A results in fewer notochord cells and therefore permits greater vacuolation per cell. By contrast, YAP inhibition during the vacuolation phase (Treatment B), when progenitor input has ceased, does not influence vacuolation dynamics. Together, these results demonstrate that reduced YAP activity in midline progenitors decreases their contribution to the notochord, and later indirectly accelerates vacuolation. Conversely, the increased packing of progenitor cells in *vgll4b* mutants (Figure 3D,E) results in a delay to anterior vacuolation, buffering the early impact of accelerated cell addition (Figure 4A,B). Taken together, we propose a mechanism by which YAP-regulated progenitor addition indirectly inhibits anterior vacuolation-mechanically coupling progenitor addition and anterior expansion across the length of the notochord.

## Discussion

In this study, we investigated the regulatory mechanisms that coordinate progenitor addition and vacuolation during zebrafish notochord morphogenesis and A-P axis elongation. Mathematical modelling and *in vivo* measurements indicated that progenitor incorporation at the posterior end contributes to tissue extension, whereas vacuolation of anterior differentiated cells generates a vacuolation front that maintains cell-size scaling along the A–P axis. YAP activity is enriched in posterior midline progenitors, coinciding with the region of active cell addition, and is negatively regulated by the inhibitor *vgll4b*. Loss of *vgll4b* results in elevated YAP activity, increased progenitor incorporation, reduced vacuolation, and, ultimately, altered notochord elongation and disrupted A–P cell-size scaling. The agreement between model predictions and experimental measurements supports the sufficiency of progenitor addition and vacuolation to explain both tissue elongation and the emergence of a linear cell-size gradient, highlighting a fundamental design principle of tissue-scale morphogenesis. These findings extend prior work on biphasic notochord elongation (Ellis et al., 2013; McLaren and Steventon, 2021) by explicitly linking progenitor dynamics with vacuolation-driven mechanical expansion.

Our data support a long-range feedback mechanism in which the balance between notochord progenitor addition and subsequent vacuolation collectively determines anterior–posterior (A–P) axis length. We propose that the effect on vacuolation is indirect: YAP primarily regulates the rate of posterior progenitor incorporation, and the resulting increase in notochord cell number subsequently attenuates anterior volume expansion driven by vacuolation that reduces notochord length. This interpretation aligns with prior zebrafish studies demonstrating that disruption of notochord vacuolation, whether by pharmacological inhibition (Tang et al., 2025; Yuan et al., 2023) or by genetic mutation affecting vacuole behaviors (Bagwell et al., 2020; Coutinho et al., 2004; Ellis et al., 2013; Garcia et al., 2017; Lim et al., 2017), leads to measurable defects in axis elongation. In *vgll4b* mutants, increased progenitor incorporation initially does not alter overall notochord length due to a buffering mechanism for natural variation in progenitor addition; however, once vacuolation becomes the dominant driver of extension, mutants display shorter notochords and reduced A–P axis length.

Our observations suggest that, during the progenitor-driven phase, the notochord retains a mechanical or morphogenetic buffering capacity that transiently compensates for elevated cell addition. This buffering fails once vacuolation dominates, revealing a feedback interaction between posterior progenitor incorporation and anterior vacuole expansion. Comparable buffering phenomena have been described in zebrafish using mechanical perturbations: robot-assisted micromanipulation has shown that axial tissues elongation resist perturbations through coordinated cell rearrangements and tension modulation (Özelçi et al., 2022). In addition, proportional regulation of posterior body elongation has been linked to spinal cord–mediated coordination of notochord and mesodermal tissue growth upon targeted ablation of spinal cord progenitors (Saunders et al., 2025). Together, these results build on prior analyses of tissue mechanics during axis elongation and highlight the notochord as a dynamic structure in which cellular composition, mechanical properties, and morphogenetic scaling converge to shape vertebrate body axes (Adams et al., 1990; Mongera et al., 2018).

Interestingly, VGLL4 has been implicated in body-size regulation in the fish *Scatophagus argus* (Yang et al., 2020), and mouse mutants exhibit a shortened body axis (Feng et al., 2019; Sheldon et al., 2022; Suo et al., 2020; Yu et al., 2019). In zebrafish, the maternal contribution of *vgll4a* and the zygotic contribution of *vgll4b* have been shown to be required for establishing the correct A–P axis length (Camacho-Macorra et al., 2024). Whereas *vgll4a* regulates YAP activity during epiboly to fine-tune axis length, the mechanism underlying the zygotic role of *vgll4b* remained unclear. Although VGLL4 is a well-established YAP inhibitor, recent studies indicate that it can also regulate gene expression independently of the VGLL4–YAP–TEAD axis in stem cell systems (Quan et al., 2023; Wang et al., 2023), mouse (Suo et al., 2025; Teng et al., 2010; Zhang et al., 2026), and in zebrafish (Fillatre et al., 2019; Lardennois et al., 2025; Wang et al., 2020; Xue et al., 2019). In our dataset, however, *vgll4b* appears to function predominantly through YAP regulation. Tailbud-specific transcriptional profiling of *vgll4b* mutants revealed upregulation of canonical YAP targets (*ccn1, ccn2a*) without significant changes in other pathways known to influence posterior progenitors. Moreover, pharmacological YAP inhibition produced phenotypes opposite to those of *vgll4b* mutants. These findings indicate that *vgll4b*’s principal role in this context is to constrain YAP activity. This spatially restricted antagonism ensures that YAP promotes progenitor maintenance and incorporation at the posterior end while preventing excessive accumulation that would compromise vacuolation.

Although our study establishes a mechanistic framework for YAP-mediated coordination of progenitor addition and vacuolation, several limitations remain. Our model represents the notochord as a one-dimensional chain of uniform cells, which may oversimplify the three-dimensional architecture as well as it’s mechanical properties. Furthermore, the downstream YAP effectors in midline progenitors and the mechanisms governing their incorporation into the notochord remain to be identified. While YAP is a prominent promoter of cell proliferation in both physiological and cancer contexts (Luo et al., 2023; Pocaterra et al., 2020), the extremely rapid pace of zebrafish development, coupled with low proliferation rates in the tailbud, indicates that axis elongation is driven primarily by cell movements and rearrangements rather than cell division (Attardi et al., 2018; Bouldin et al., 2014; Kanki and Ho, 1997). This is consistent with observations that pharmacological inhibition of proliferation does not prevent A–P axis extension (Stooke-Vaughan et al., 2025). Notably, VGLL4 antagonism of YAP activity has been shown to reduce cell migration in cultured cells (Mickle et al., 2021; Sun et al., 2022) and *in vivo* models (Zhang et al., 2014; Zhang et al., 2017). We therefore hypothesize that elevated YAP activity in *vgll4b* mutant progenitors increases their rate of incorporation into the notochord rather than enhancing their proliferation.

In summary, our study identifies YAP as a key regulator of progenitor addition and vacuolation during zebrafish notochord morphogenesis. By integrating experimental data with computational modelling, we demonstrate that a feedback mechanism balances posterior progenitor incorporation with anterior vacuole expansion to maintain notochord elongation and A–P scaling. Understanding how molecular signals interface with tissue mechanics to ensure robust scaling has broad implications for congenital axial malformations and may inform strategies for engineering notochordal or axial tissues (Nájera et al., 2025). More broadly, the interplay between progenitor dynamics and volumetric expansion may represent a general principle through which embryos coordinate multiple morphogenetic processes to achieve robust patterning and size control.

**Supplementary Figure 1.**
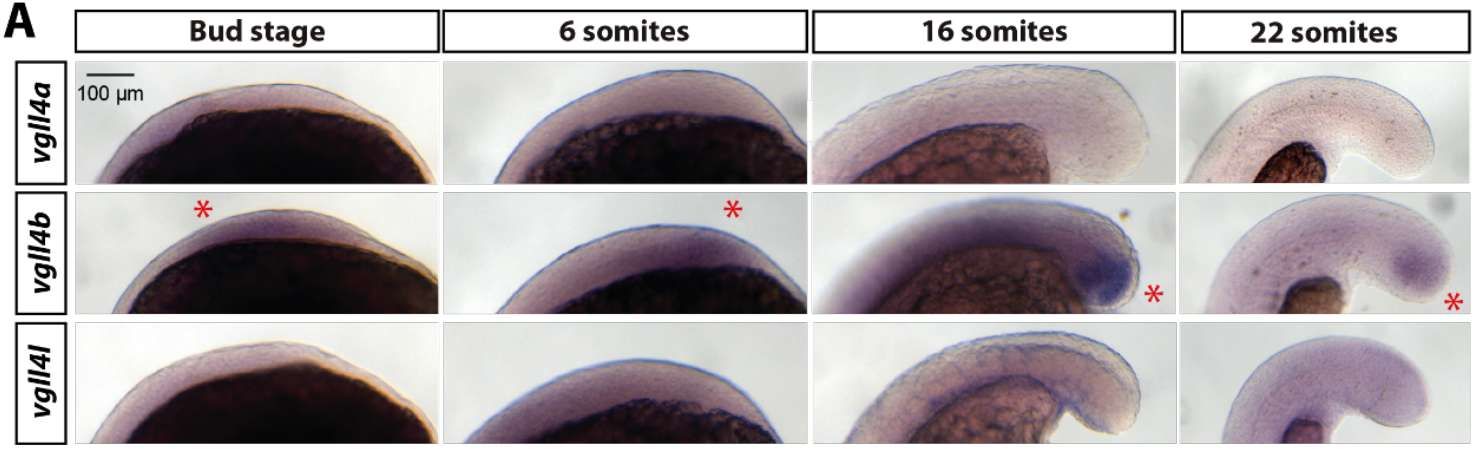
Expression patterns of zebrafish *vgll4* paralogues in the posterior embryonic axis during A–P elongation. **A**. ISH staining of VGLL4 zebrafish paralogues *vgll4a, vgll4b*, and *vgll4l* in the tail at different developmental stages (scale bar: 100 μm). Red asterisk points *vgll4b* expression.

**Supplementary Figure 2.**
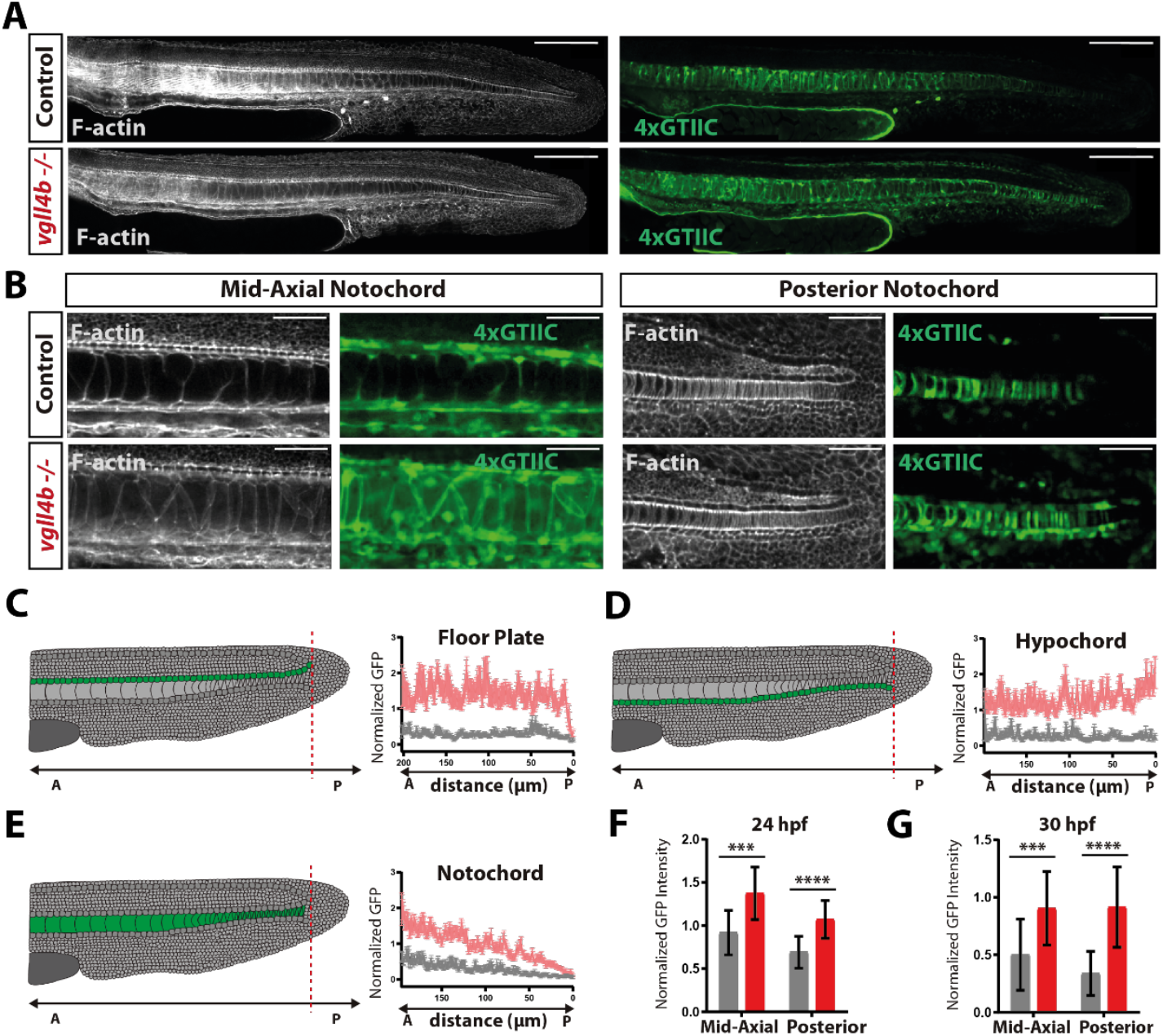
Increased YAP activity in the midline structures of *vgll4b* mutants. **A**. Confocal images of 30 hpf *Tg(4xGTIIC:GFP)* control and *Tg(4xGTIIC:GFP)*; *vgll4b*−/− embryos (scale bar: 100 μm). **B**. Confocal images of the mid-axial and posterior notochord in control and mutant embryos at 30 hpf (scale bar: 50 μm). **C, D, E**. Quantification of normalized GFP intensity across the floor plate, notochord, and hypochord at 30 hpf, showing sustained YAP overactivation in mutants (control n = 17, mutant n = 23), the control group is represented by the gray line, and mutants by the red line. **F**. Quantification of normalized GFP intensity in the mid-axial and posterior notochord at 24 hpf: mid-axial (t-test; control n = 10, mutant n = 18; p = 0.0005), posterior (t-test; control n = 13, mutant n = 17; p < 0.0001). **G**. Quantification of normalized GFP intensity in the mid-axial and posterior notochord at 30 hpf: mid-axial (t-test; control n = 16, mutant n = 21; p = 0.0005), posterior (Mann–Whitney test; control n = 18, mutant n = 23; p < 0.0001).

**Supplementary Figure 3.**
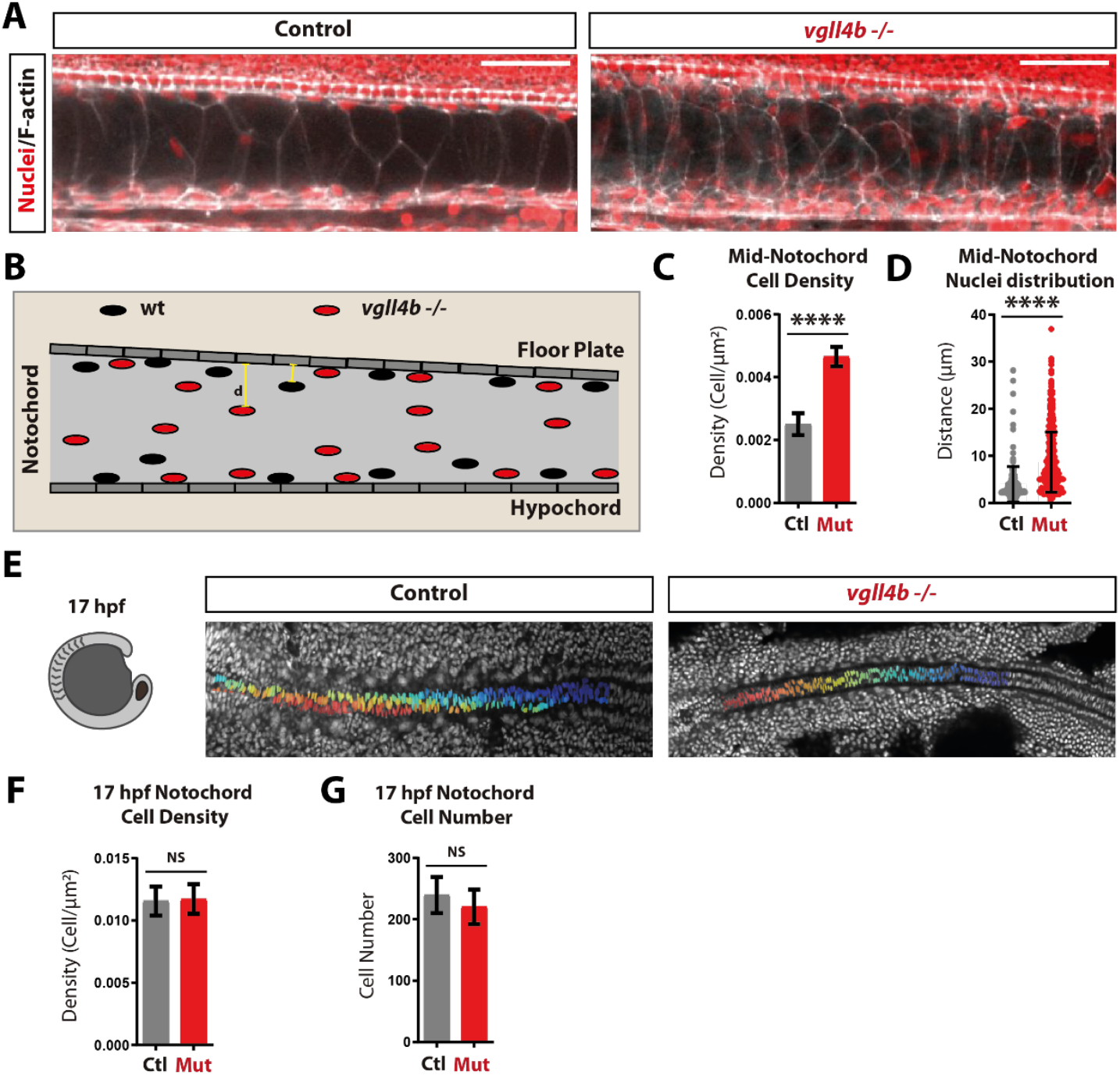
Elevated notochord cell density in *vgll4b* mutants alters the spatial distribution of nuclei within the notochord. **A**. Confocal images of the mid-axial notochord in control and mutant embryos at 30hpf stained with DAPI and phalloidin (scale bar: 50 μm). **B**. Schematic illustrating nuclear distribution in mid-axial notochord cells of control and mutant embryos. **C**. Quantification of mid-axial notochord cell density (t-test; control n = 5, mutant n = 5; p < 0.0001). **D**. Quantification of mid-axial notochord nuclear distribution (Mann–Whitney test; control n = 5, mutant n = 5; p < 0.0001). **E**. Confocal images of the notochord in control and mutant embryos at 17 hpf, prior to tail formation. Notochord nuclei were segmented for quantitative analysis. **F**. Quantification of notochord cell density at 17 hpf (t-test; control n = 10, mutant n = 7; p = 0.7926). **G**. Quantification of notochord cell number at 17 hpf (t-test; control n = 10, mutant n = 7; p = 0.1981).

**Supplementary Figure 4.**
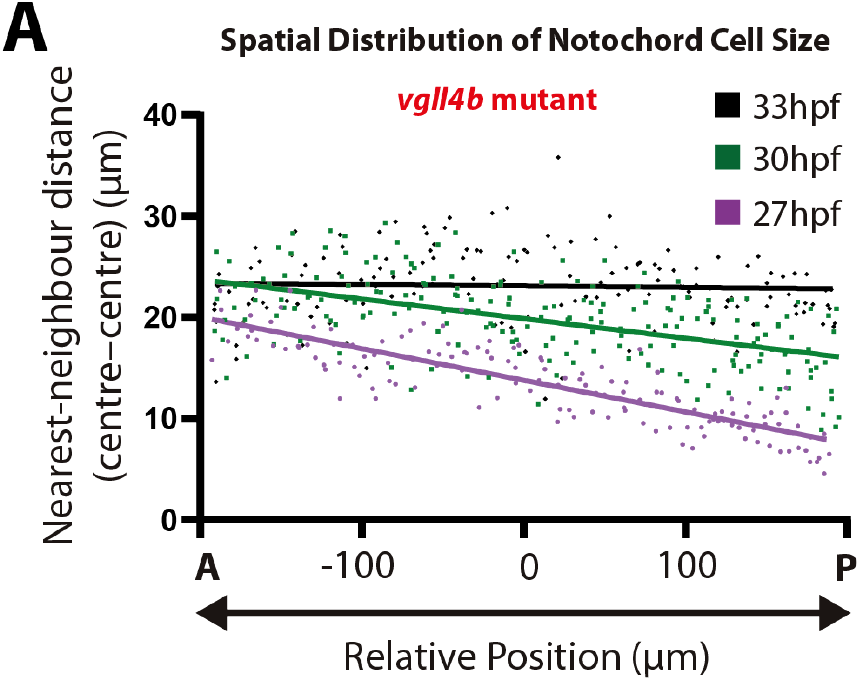
Altered Spatial Distribution of Notochord Nearest Neighbour Cell Distances centre to centre in *vgll4b* Mutant Embryos. **A**. Quantification of AP profiles of nearest-neighbour cell distances centre to centre within the notochord at 27 hpf, 30 hpf, and 33 hpf in *vgll4b* mutant embryos. The A– P axis in the graph is centered at the yolk-level extension. The region of interest (ROI) used for measurements spanned 200 µm on each side of this reference point, for a total length of 400 µm.

**Supplementary Figure 5.**
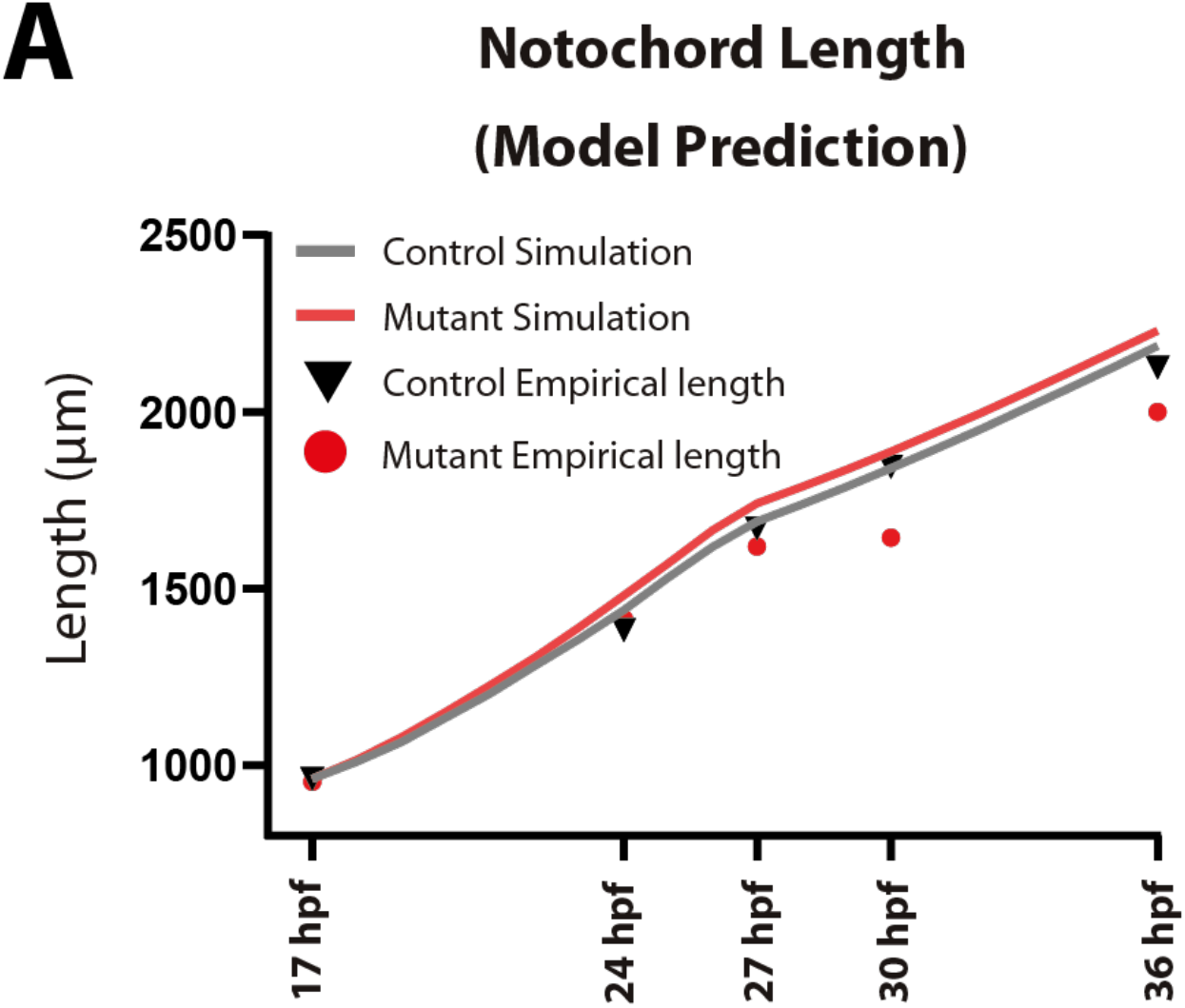
Predictions of mathematical model of notochord length. **A**. Mathematical model predictions of notochord length with vacuolation rate *J* and progenitor addition rate *r*_*p*_ as functions of normalized YAP levels *Y* for control and *vgll4b* mutant embryos at different time points (see Eqs. 1 and 2 in the main text). The model assumes that YAP does not affect the vacuolation front speed *v*_front_. The empirical measurements shown in the graph are the same as those presented in Figure 4A. The parameters used for the simulations are the same as the one used in Table 3 except normalized wild-type Yap level (0.2), normalized mutant Yap level (0.23), the maximum rate of vacuolation-induced cell growth 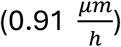 and the maximum progenitor addition rate 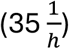.

**Supplementary Figure 6.**
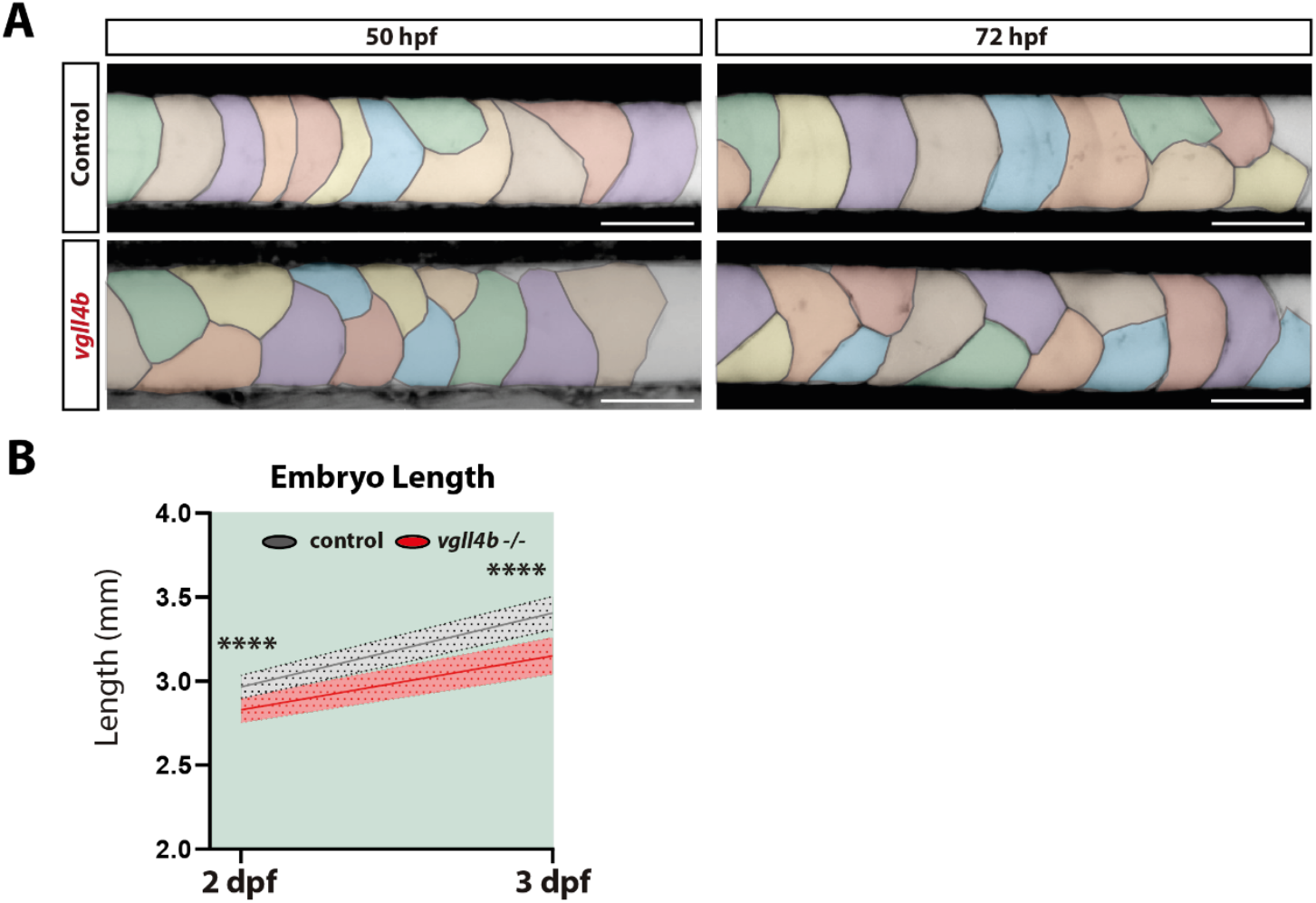
Reduced axial length in *vgll4b* mutant embryos at later developmental stages. **A**. Segmentation of notochord vacuoles in control and mutant embryos at 50 hpf and 72 hpf visualized with Bodipy staining (scale bar: 50 μm). **B**. Quantification of embryo length at 2 dpf (t-test; control n = 49, mutant n = 58; p < 0.0001) and 3 dpf (t-test; control n = 135, mutant n = 147; p < 0.0001).

**Supplementary Figure 7.**
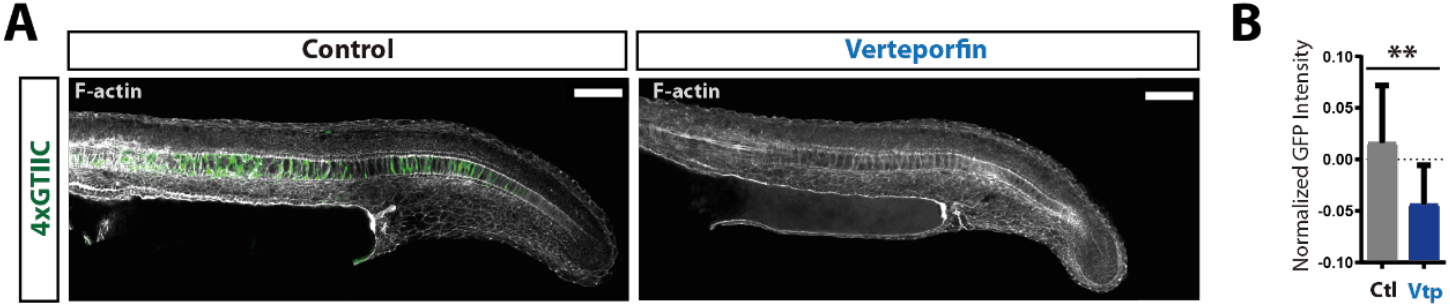
Verteporfin treatment inhibits GFP expression in *Tg(4xGTIIC:GFP)* embryos. **A**. Confocal images of 24 hpf control and verteporfin-treated *Tg(4xGTIIC:GFP)* embryos (scale bar: 100 μm). **B**. Quantification of normalized GFP intensity in the mid-axial notochord, showing significant reduction in treated embryos (t-test; control n = 11, treated n = 16; p = 0.0034).

## Materials and Methods

### 1D model of zebrafish notochord elongation

We modelled the notochord as a 1D tissue composed of cells. Each cell is represented as a spring–mass element: a spring with elastic constant *k* anchored at one end and connected to a mass *m* at the other. The tissue consists of a sequence of these spring–mass units arranged along the *x*-axis, corresponding to the anterior–posterior (AP) axis. The first spring is attached to a fixed anchor point (a ‘wall’) at the most anterior end of the notochord. New cells are added posteriorly by attaching the anchor point of the new cell to the mass of the current last cell.

All cells obey Hooke’s law with the same elastic constant k, the same mass *m* (set arbitrarily to 1), and a target length *S*_*i*_, defined as the spring’s rest length (see Figure 1B for a schematic). The position of mass *i* (and thus of cell *i*) changes according to the net force exerted by its two neighbouring springs, 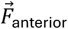 and 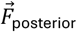. A viscous drag force, appropriate for the low–Reynolds number regime, opposes motion. We assumed that notochord cells are in an overdamped regime where the inertial term is negligible compared with the viscous forces. Applying Newton’s second law for the mass *i* gives:

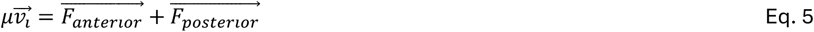

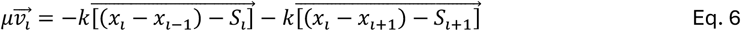

where 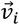 is the velocity of cell *i, x*_*i*_ is its position, *S*_*i*_ is its target size, and *μ* is the drag coefficient.

We assumed the presence of a vacuolation front that starts at time *t* = *t*_start_vac_ at *x* = 0 (the anterior anchor point) and moves posteriorly at constant speed *v*_front_. A cell becomes vacuolated when its position lies anterior to the front. Upon vacuolation, the target size *S*_*i*_ increases according to

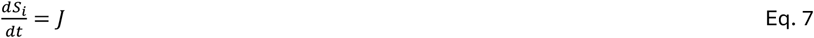

where *J* is a constant representing the vacuolation-induced rate of size increase. The solution is

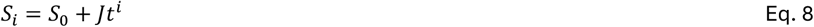

where *S*_0_ is the initial cell size (identical for all cells), and *t*^*i*^ = *t* − *t*^*vacuolated*^ is the time elapsed since cell *i* became vacuolated.

Progenitor addition is implemented by inserting new cells at the posterior end at regular intervals *t*_add_, related to the progenitor addition rate *r*_*p*_ by *r*_*p*_ = 1/*t*_add_. Progenitor addition stops after a time *T*_*p*_ from the start of the simulation. Each time a progenitor cell is added, a small anteriorly directed force is applied to its mass, effectively compressing the tissue. In the initial condition of the simulation, we assumed *N*_0_ cells of size *S*_0_ distributed between *x* = 0 and *x* = *L*_0_, representing the anterior and posterior borders of the notochord, respectively. This initial configuration corresponds to the notochord at 16 hpf, with the vacuolation wave initiating at *x* = 0 while a progenitor is added at *x* = *L*_0_. All model parameters are depicted in Table 3.

**Table 1.**
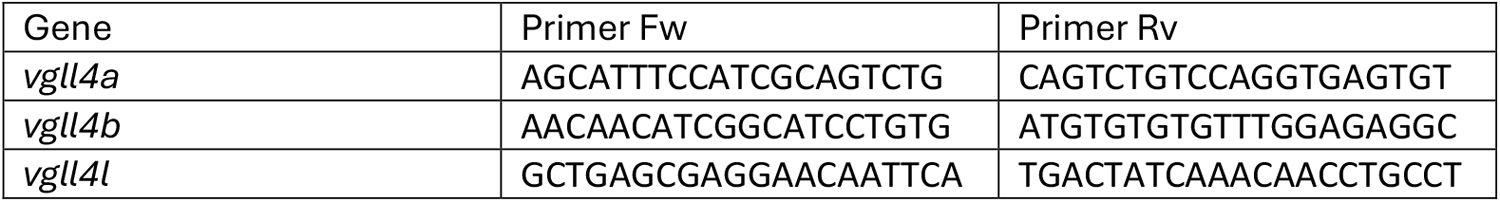
Primers used for ISH probe generation.

**Table 2.**
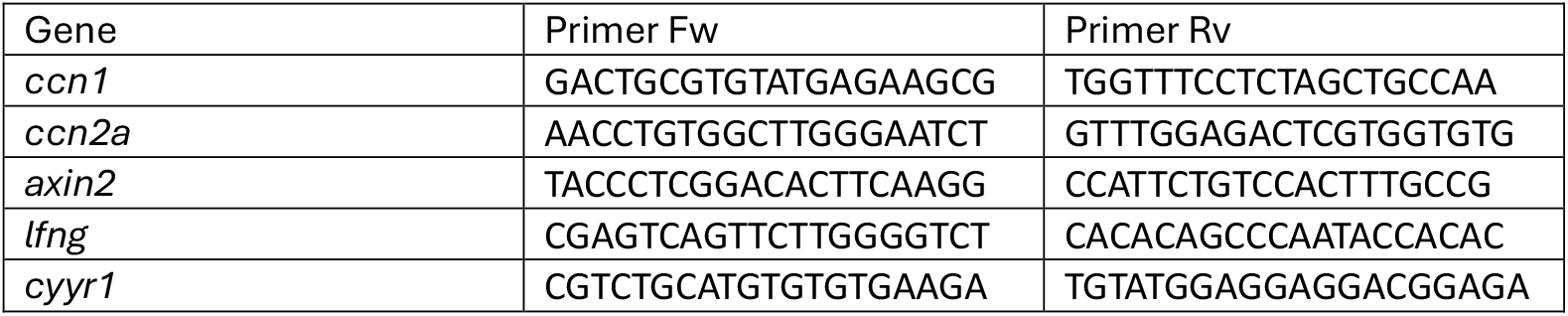
Primers used for qPCR experiments.

**Table 3.**
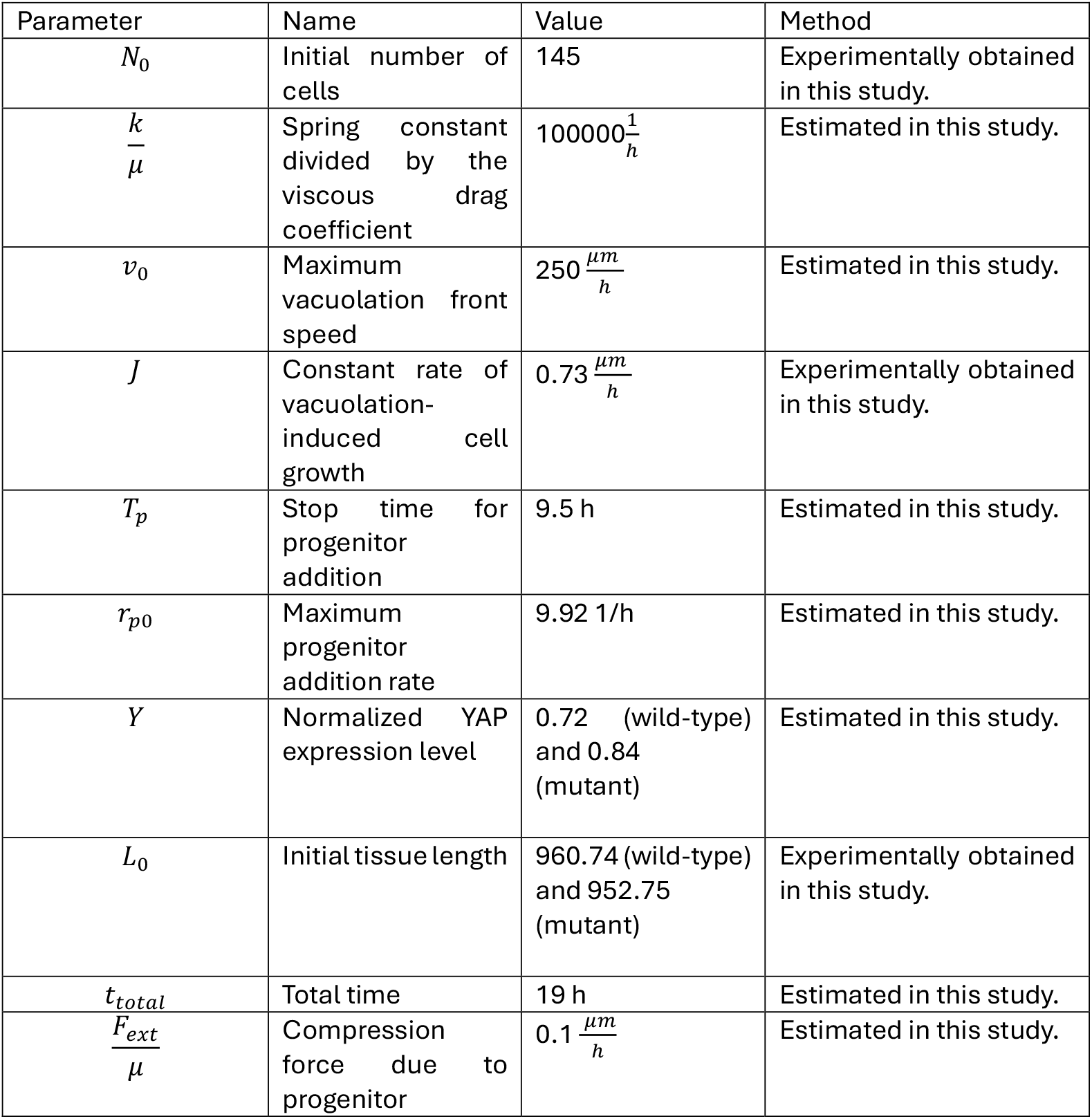
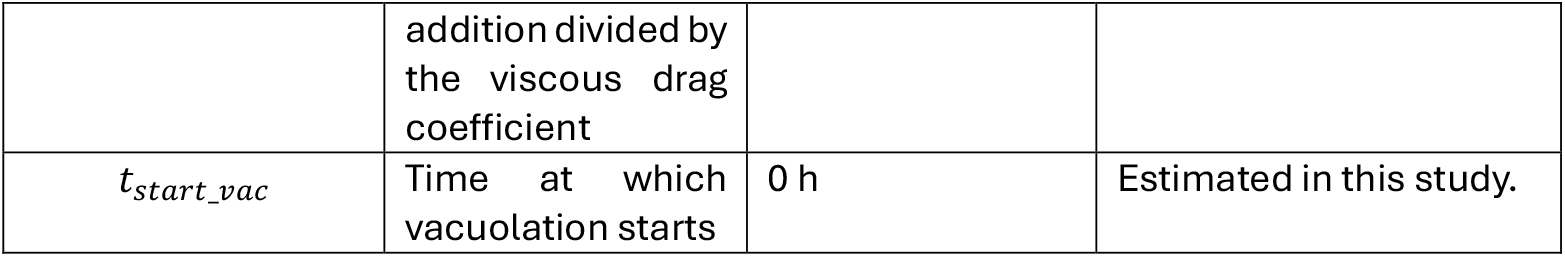
Parameters corresponding to the mathematical model described by Eqs. 3 and 4 in the main text.

| Parameter | Name | Value | Method |
| --- | --- | --- | --- |
| $N_0$ | Initial number of cells | 145 | Experimentally obtained in this study. |
| $\frac{k}{\mu}$ | Spring constant divided by the viscous drag coefficient | $100000 \frac{1}{h}$ | Estimated in this study. |
| $v_0$ | Maximum vacuolation front speed | $250 \frac{\mu m}{h}$ | Estimated in this study. |
| $J$ | Constant rate of vacuolation-induced cell growth | $0.73 \frac{\mu m}{h}$ | Experimentally obtained in this study. |
| $T_p$ | Stop time for progenitor addition | 9.5 h | Estimated in this study. |
| $r_{p0}$ | Maximum progenitor addition rate | 9.92 1/h | Estimated in this study. |
| $Y$ | Normalized YAP expression level | 0.72 (wild-type) and 0.84 (mutant) | Estimated in this study. |
| $L_0$ | Initial tissue length | 960.74 (wild-type) and 952.75 (mutant) | Experimentally obtained in this study. |
| $t_{total}$ | Total time | 19 h | Estimated in this study. |
| $\frac{F_{ext}}{\mu}$ | Compression force due to progenitor addition divided by<br>the viscous drag<br>coefficient | $0.1 \frac{\mu m}{h}$ | Estimated in this study. |
| $t_{start\_vac}$ | Time at which<br>vacuolation starts | 0 h | Estimated in this study. |

### Condition for cell-size scaling in the notochord

In this section, we show that during the second phase, after progenitor addition ceases, the model predicts a linearly decreasing cell-size profile along the AP axis. We refer to this phenotype as *scaling*, because the shape of the distribution is conserved as the tissue grows.

This behaviour arises because vacuolated cells are pushed posteriorly by the expansion of anterior cells. Specifically, when a cell *i* at position *x*_*i*_ becomes vacuolated, its posterior displacement is driven by the *N*(*x*_*i*_, *t*) anterior cells that are also enlarging. Although the position of cell *i* could in principle be determined from the positions of all anterior cells, its velocity can be expressed more directly: it equals the cumulative size-increase rate of all anterior cells.

We assume that all cells are modelled as rigid springs (with high spring constant *k*), so that their actual length closely follows their target length *S*_*i*_; hence, we refer to cell length and target length interchangeably. Since there are *N*(*x*_*i*_, *t*) anterior cells and each grows at rate *J*, the posterior velocity of cell *i* is given by *v*_*i*_ = *N*(*x*_*i*_, *t*)*J*.

From the perspective of a reference frame centred on cell *i*, the vacuolation front appears to move with an effective speed

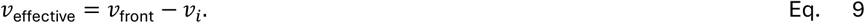

In this reference frame, the vacuolation wave front therefore advances at a constant speed. From the time the cell became vacuolated, 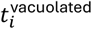, up to the current absolute time *t*, the front has travelled a distance Δ*x*. This distance corresponds to the separation between the front and the cell:

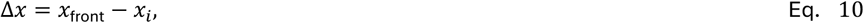

where *x*_*i*_is the position of cell *i*. Because the vacuolation wave front moves at constant speed in this cell-centred frame, we can also express this distance as

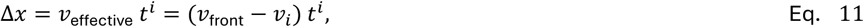

where 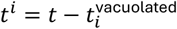 is the time elapsed since the front passed cell *i*.

Combining the two expressions for Δ*x* gives

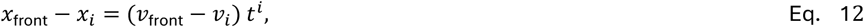

and therefore

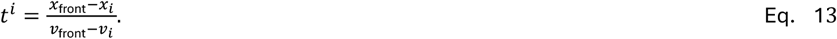

Finally, since the vacuolation wave front advances at constant speed, its position is simply

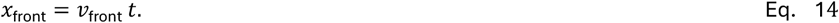

Hence, we obtain:

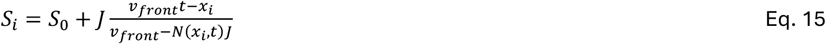

We can replace the index *i* by a general position *x*, yielding:

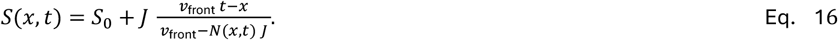

If *v*_front_ ≫ *N*(*x, t*) *J*, this expression simplifies to:

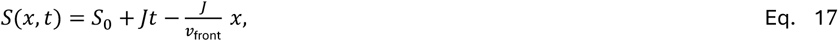

which is a straight line in *x* with constant slope and an intercept that increases linearly in time:

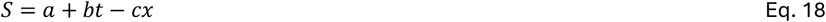

Where a = *S*, b = *J* and 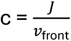. We refer to this as scaling when the spatial distribution of cell sizes at different times forms straight lines with identical slopes but different intercepts.

To study under what conditions this linear profile of cell sizes emerges, we bounded the term *N*(*x, t*) in Eq. 14. Because new cells are added exclusively at the posterior tip, *N*(*x, t*) must lie between the initial number of cells and the maximum number that can accumulate during the period of progenitor addition. Thus, *N*(*x, t*) is bounded as follows:

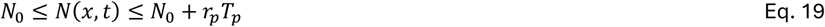

The condition *v*_front_ ≫ *N*(*x, t*) *J* is thus guaranteed for all *x* and *t* provided that

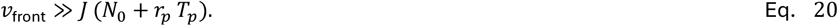

We define the dimensionless parameter

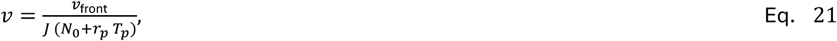

which represents the ratio between the speed of the vacuolation front and the maximal speed at which the notochord could expand.

The condition for obtaining the linear profile *S* = *a* + *bt* − *cx* therefore simply becomes

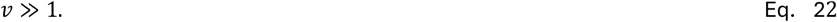

When *v* ≫ 1, we can ensure that the spatial distribution of cell sizes along the notochord is a straight line with a constant slope and a time-increasing intercept.

### Constraining the model parameter space

Eq. 17 describes a linear function for the spatiotemporal distribution of cell notochord sizes under the hypothesis that *v*_front_ ≫ *N*(*x, t*) *J*. If we look at this equation at a fixed time, we can realize that the slope of it is 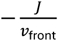. Alternatively, if we look at it at a fixed position, the slope is now *J*. If we measure these two slopes, we will end up with two equations with two unknowns (*J* and *v*_front_) which we can solve for.

We fitted a line at the AP profiles of nearest-neighbour cell distances centre to centre within the notochord at 27, 30 and 33 hpf for the wild-type (Fig.1H) and mutant (Sup. Fig 4A). To calculate the slope of eq. 17 at fixed time, we estimated the average and standard deviation of the slopes of the fitted lines at 27, 30 and 33 hpf for the wild-type and we repeated this procedure for the mutant. To estimate the slope of the wild-type at fixed position, we followed this procedure. We selected a position and calculated the cell size of the wild-type at that position at 27, 30 and 33 hpf from their corresponding fitted lines. We took those three values and fitted a line to AP profiles of nearest-neighbour cell distances centre to centre within the notochord as a function of time and we obtained the slope of that line. Then we repeated this procedure with other positions within the measured domain spaced by 0.001 µm and calculated the slope again. Finally, we calculated the mean and standard deviation among all those slopes. We repeated this procedure with the mutant. With the two slopes estimated, we solve the system of equations to calculate the values of *J* and *v*_front_ and used propagation of errors to calculate the standard deviation of *v*_front_.

### Animal husbandry

Adult zebrafish were maintained and all regulated procedures were conducted in accordance with the Animals (Scientific Procedures) Act 1986, as amended in 2012, following approval by the Animal Welfare and Ethical Review Body (AWERB) at the University of Cambridge. Embryos were kept in standard E3 medium throughout all experimental procedures. Developmental staging followed established criteria (Kimmel et al., 1995). To suppress spontaneous muscle contractions, embryos were incubated in tricaine (ethyl 3-aminobenzoate; Sigma-Aldrich, A5040) at a final concentration of 0.16 g/mL in E3 medium. The zebrafish lines used were Tg(4xGTIIC:GFP) (Miesfeld and Link, 2014) and *vgll4b*-P156Rfs5 (Camacho-Macorra et al., 2024).

### Pharmacological treatments

Live imaging was performed using the vital dye BODIPY TR methyl ester (MED; Invitrogen), following standard protocol (Ellis et al., 2013). Embryos were incubated in 2% MED for 45 min, rinsed several times in E3 medium, mounted, and imaged immediately.

For verteporfin treatment, dechorionated embryos were maintained at 28 °C in E3 medium containing either DMSO (vehicle control) or verteporfin (Tocris, 5305) dissolved in DMSO. Verteporfin was applied at a final concentration of 10 μM.

### Fixation and staining

Embryos were fixed in 4% paraformaldehyde for 24 h at 4 °C. After fixation, samples were washed in phosphate-buffered saline without calcium and magnesium [PBS(–/–)] containing 0.05% Tween-20 (PBST). Nuclei were stained with DAPI (1:500), and filamentous actin was labeled with Alexa Fluor 647–conjugated phalloidin (1:1000) in PBST supplemented with 0.1% DMSO. Staining was carried out for at least 24 h at 4 °C.

ISH probes (Supplementary Figure 1) specific for each VGLL4B paralogue (*vgll4a, vgll4b*, and *vgll4l*) were generated by RT-PCR using cDNA from different developmental stages and gene-specific primers (Supplementary Table 1) with the Expand High Fidelity PCR System (Roche). Amplicons were cloned using the StrataClone PCR Cloning Kit (Agilent). Digoxigenin-UTP–labelled antisense riboprobes were synthesized in vitro using the DIG RNA Labeling Mix (Roche). Probes were precipitated overnight in LiCl at –20 °C, pelleted by centrifugation at 12,000 × g for 30 min at 4 °C, washed with 70% ethanol, centrifuged again at 12,000 × g for 15 min, dried at 65 °C, and resuspended in a 1:1 mixture of RNase-free water and formamide (ITW reagents). ISH was performed using standard procedures and visualized with NBT/BCIP.

Fluorescence in situ hybridization was performed using the hybridization chain reaction method (Choi et al., 2018). Immunohistochemistry followed (Sorrells et al., 2013). Primary antibodies were chicken anti-GFP (1:200; Abcam, Ab13970; RRID:AB_300798) and rabbit monoclonal anti-YAP (1:1000; Cell Signaling Technology, 8418). Secondary antibodies (all goat-raised; 1:1000) were Alexa Fluor 488 anti-chicken (Thermo Fisher Scientific, A-11039) and Alexa Fluor 633 anti-rabbit (Invitrogen, A21071).

### Confocal microscopy

For live imaging, embryos were mounted as described (Hirsinger and Steventon, 2017). For fixed samples, embryos were fully dissected away from the yolk, and the head was removed to allow the body to lie flat along the lateral axis. Embryos were mounted in 80% glycerol between two 1H coverslips adhered with double-sided tape. Imaging was performed on an inverted Zeiss LSM700 confocal microscope. Whole embryos were imaged in tiled sections using a 20× objective with 2× line averaging. Laser intensities, gain, and pixel resolution were kept constant across experiments.

### Photolabelling

Photolabelling was performed using the photoconvertible protein Kikume (Hatta et al., 2006). The plasmid nls-Kikume (gift from Ben Martin, SUNY Stony Brook, USA) was used to synthesize mRNA with the mMessage mMachine kit (Thermo Fisher Scientific, AM1344). Embryos were microinjected prior to the first cell division with a 100 μm diameter doplet containing 300 ng/μL Kikume mRNA. Photoconversion was performed using a 405 nm laser on a Zeiss LSM700 confocal microscope. A 30-second exposure was applied to a rectangular region of interest (ROI) positioned at the posterior end of the notochord. Upon exposure, the Kikume fluorophore underwent a shift from green fluorescence (KikG) to red fluorescence (KikR), indicating successful photoconversion.

### Visualization and registration

Image analysis was performed using Fiji (Schindelin et al., 2012). Two-dimensional length measurements were obtained using Fiji’s segmented line region of interest (ROI) tool. Individual image tiles were acquired and subsequently stitched using a custom script that performed successive rounds of pairwise stitching via the Fiji Stitching plugin (Preibisch et al., 2009). When necessary, images were rotated using the TransformJ plugin (Meijering et al., 2001) to align anatomical structures.

### Measurements and quantifications

Tissue lengths were measured using Fiji’s segmented-line tool. Notochord length was defined from the anterior boundary of the first somite to the posterior end of the notochord.

Notochord vacuole area was quantified from the central plane of Z-stacks. Individual vacuoles in BODIPY-stained embryos were manually segmented, and their areas measured. To determine the position of each cell along the axis, a 400 μm rectangular ROI centered on the yolk extension was used. Each vacuole within the ROI was segmented, polygon coordinates were extracted, and centroids calculated.

Cell density at the posterior notochord was quantified by selecting the central plane of the Z-stack and applying a 100 μm ROI positioned using the floor plate and hypochord as landmarks. DAPI-labelled nuclei were counted and normalized to the ROI area. For notochord cell quantification at 17 hpf (Supplementary Figure 3E), a region of interest (ROI) encompassing the notochord between the anterior boundary of the first somite and the posterior boundary of the last somite was defined. Nuclei within this ROI were subsequently segmented using Cellpose software.

To assess nuclear distribution at the mid-notochord in control and mutant embryos, a 400 μm square ROI centered on the yolk extension was applied. The distance from each DAPI-labelled nucleus to the nearest boundary (floor plate or hypochord) was measured using the segmented-line tool.

For GFP intensity quantification (Figure 2F–G; Supplementary Figure 5A), a 400 μm ROI centered on the yolk extension in the central plane of the notochord was used. GFP intensity was normalized to phalloidin intensity within the same ROI. To quantify GFP along the posterior floor plate, hypochord, or notochord, a straight line was drawn from the posterior notochord anteriorly, and pixel intensities extracted. GFP values were normalized to phalloidin and corrected by subtracting background signal measured from a separate region where no GFP was expressed.

Volume quantification of expression domains was performed using Imaris. Surfaces of interest were segmented from confocal stacks, and volumes calculated. Each volume was normalized to the total volume of the DAPI-labelled tailbud, defined using the posterior boundary of the last somite.

For measurements of embryo length at 2 and 3 dpf, embryos were anesthetized in tricaine, oriented vertically, and imaged at 13X magnification using a Leica MZ10F stereomicroscope equipped with a Leica 295 camera.

### qPCR Experiments

Total RNA was extracted with TRIzol (Thermo Fisher Scientific) and resuspended in nuclease-free water. RNA concentration was quantified using the Qubit RNA High Sensitivity Kit (Thermo Fisher Scientific, Q32852). First-strand cDNA was synthesized using the SuperScript III First-Strand Synthesis System (Thermo Fisher Scientific, 18080051). For primer standardization, control cDNA was diluted 1:10, 1:20, 1:30, and 1:40. Ct values for each gene-specific primer pair were compared with those of the housekeeping gene cyyr1 (Casadei et al., 2011). Standardized primers were then used to quantify gene expression across treatment groups. All experimental qPCRs were performed using cDNA diluted 1:40. Primer sequences are listed in Supplementary Table 2.

### Statistical analysis

All graphs and statistical analyses were generated using GraphPad Prism 7. Two-group comparisons were performed using unpaired t-tests for data with parametric distributions or Mann–Whitney tests for data with non-parametric distributions. Statistical significance was reported using the following thresholds: *p* < 0.05 (\**), p < 0.01 (**), p < 0.001 (\*\*\**), and *p* < 0.0001 (****).

## ACKNOWLEDGEMENTS

We thank Professor Paola Bovolenta for her support, as the *vgll4b* mutant fish line was generated by the first author in her laboratory at the Centro de Biología Molecular Severo Ochoa (CBMSO), Spain. We thank Yuri Takahashi, Alice C. Yuen, Qiyu Chen, and Ye Zhao for reviewing and commenting on the manuscript. B.J.S. and C.C-M are supported by a Wellcome Trust Discovery Award (225360_Z_22_Z). O.C. and A.C.S were funded by Fondo para la Investigación Científica y Tecnológica (PICT-2019-03828) and the BBSRC grant BB/X014908/1, both to O.C.

